# Novel Universal Primers for Metabarcoding eDNA Surveys of Marine Mammals and Other Marine Vertebrates

**DOI:** 10.1101/759746

**Authors:** Elena Valsecchi, Jonas Bylemans, Simon J. Goodman, Roberto Lombardi, Ian Carr, Laura Castellano, Andrea Galimberti, Paolo Galli

## Abstract

Metabarcoding studies using environmental DNA (eDNA) and high throughput sequencing (HTS) are rapidly becoming an important tool for assessing and monitoring marine biodiversity, detecting invasive species, and supporting basic ecological research. Several barcode loci targeting teleost fish and elasmobranchs have previously been developed, but to date primer sets focusing on other marine megafauna, such as marine mammals have received less attention. Similarly, there have been few attempts to identify potentially ‘universal’ barcode loci which may be informative across multiple marine vertebrate Orders. Here we describe the design and validation of four new sets of primers targeting hypervariable regions of the vertebrate mitochondrial 12S and 16S rRNA genes, which have conserved priming sites across virtually all cetaceans, pinnipeds, elasmobranchs, boney fish, sea turtles and birds, and amplify fragments with consistently high levels of taxonomically diagnostic sequence variation. ‘*In silico*’ validation using the OBITOOLS software showed our new barcode loci outperformed most existing vertebrate barcode loci for taxon detection and resolution. We also evaluated sequence diversity and taxonomic resolution of the new barcode loci in 680 complete marine mammal mitochondrial genomes demonstrating that they are effective at resolving amplicons for most taxa to the species level. Finally, we evaluated the performance of the primer sets with eDNA samples from aquarium communities with known species composition. These new primers will potentially allow surveys of complete marine vertebrate communities in single HTS metabarcoding assessments, simplifying workflows, reducing costs, and increasing accessibility to a wider range of investigators.

## INTRODUCTION

The use of DNA fragments extracted from environmental sources (e.g. soil and water samples) is becoming a well-established tool for monitoring biodiversity (Deiner et al. 2017; Jarman et al. 2018). Within the marine environment, such eDNA surveys have been used to assess the diet of marine species (Deagle et al. 2010, Peters et al.2015, McInnes et al. 2017), monitor the species diversity of marine communities (Port et al. 2016; Sigsgaard et al. 2017), determine the presence/absence of invasive species (Borrell et al. 2017) and to obtain estimates of population genetic diversity (Sigsgaard et al. 2016). Community biodiversity surveys from eDNA (i.e. eDNA metabarcoding) rely on primers targeting specific taxonomic groups and high-throughput sequencing to amplify and sequence barcoding regions from all species of interest (Creer et al. 2016). While DNA metabarcoding primers have been developed to target several individual marine taxonomic groups (e.g. elasmobranchs, teleost fish, cephalopods and crustaceans) (e.g. Jarman et al. 2016, Miya et al. 2015, Komai et al. 2019, Bylemans et al. 2018), to date no primer sets have been specifically designed to maximize the recovery and identification of marine mammals.

The few marine eDNA studies focussing on marine mammals used targeted species-specific assays (Foote et al. 2012, Baker et al. 2018) or used universal fish specific primers as a proxy to assess the total vertebrate biodiversity (e.g. Andruszkiewicz et al. 2017). All these approaches present at least one drawback when aiming to detecting the presence of cetacean and pinniped species within an eDNA sample. Primer sets designed for one group, i.e. ‘fish-specific’ primers, might amplify eDNA from other taxa, but unquantified primer mismatches risk reduced detection rates and the introduction of biases in high throughput sequencing (HTS) results (e.g. Elbrecht et al 2018). Where primers are designed for the species of interest, amplicon target size may also be a consideration. The instability and short life span of eDNA molecules (e.g. Thomsen et al. 2012), favours the use of short and informative (hypervariable) DNA regions, such as the mitochondrial 12S, 16S and cytochrome oxidase I (*COI*) genes, with amplicons of typically around 100-200 bp although larger mtDNA fragments have been shown to be successfully amplified from eDNA samples (Deiner et al. 2017).

There is therefore a need to develop marine mammal-specific primers suitable for metabarcoding analysis from marine eDNA samples. Mitochondrial 12S and 16S regions provide suitable targets since their sequence variation provides good taxonomic resolution for macro-eukaryotes, while also maintaining conserved sites across regions for siting primers (Deagle et al. 2014). The design criteria for such primers should be, a) primer sites which are conserved among marine mammal groups, while amplifying hypervariable DNA fragments for taxonomic resolution; b) where possible, identify marine mammal specific priming sites in order to reduce cross-amplification with human DNA, thus lessening contamination risks from investigators, swimmers, or other biological residues left by humans; c) for each primer set evaluate predicted binding efficiency and amplicon sequence diversity for other marine vertebrates such as fish and sea turtles (when necessary allowing for a single degenerate base per primer). This final point would provide to give a more accurate understanding of primer specificity and suitability for use with other vertebrate groups. Primer sets proven to have reliable affinity across multiple vertebrate Orders and could support more efficient and cost-effective eDNA biodiversity surveys.

In this paper we report on the development of novel “universal” marine vertebrate eDNA primers meeting the above criteria, their *‘in silico*’ validation against a large marine vertebrate NCBI-Genbank dataset, and their successful initial application and validation with a high throughput sequencing analysis of environmental water samples collected from a public marine aquarium. Finally, we test for inter- and intra-specific variation over large mitogenomic data sets available for marine mammal species, presenting ready-to-use guidelines for the accurate selection of primer set of choice in specific marine mammal studies.

## MATERIALS AND METHODS

### Initial design of primer sets

Seventy-one complete mitochondrial genome sequences, representative of most marine vertebrate groups (fish, sea turtles, birds and marine mammals), were retrieved from GenBank and used for initial primer development (Online Resource 1 in Supplementary Materials). The selected sequences represented 30 marine vertebrate families, including most marine mammal families (all 3 Pinniped families, both Sirenian and 13 Cetacean families). The selection comprised all Cetacean species occurring in the Mediterranean Sea. In addition, four human mitochondrial genomes representative of the four main human haplogroups (i.e. haplotypes 16, 31, 33 and 52 in Ingman et al. 2000) were included (Online Resource 1) in order to design primers with reduced amplification efficiency for human DNA. All sequences (n = 75) were aligned with the online tool Clustal-Omega (https://www.ebi.ac.uk/Tools/msa/clustalo/) with default parameters, and the complete ribosomal 12S and 16S genes were isolated. Potential sites for metabarcoding primers were identified by manually searching for suitable locations within alignments. Gene regions were considered suitable for designing metabarcoding primers if they encompassed a short (80-230bp) highly variable fragment, required for species discrimination, and were flanked by highly conserved sites for situating primers. Where possible, (i.e. when enough intra-mammal variation was found in proximity of the priming sites), we also tried to design, for each candidate locus, we also tried to design alternative primers which minimised the probability of amplifying human targets, by ensuring mismatches between the primers and human templates. Such variants could be preferentially used in studies specifically targeting marine mammals.

### Primer evaluation and validation

Three approaches were used to assess primer performance. Firstly, primers were evaluated *in silico* in two steps: i) predicted primer binding and amplicon sequence diversity was assessed using the ecoPCR scripts within the OBITOOLS software package (Ficetola et al. 2010; Boyer et al. 2016); and ii) 680 complete marine mammal mitogenome sequences deposited in Genbank, were used to quantify sequence diversity for the primer target regions within marine mammal Families, Genera and species, to provide recommendations on taxonomic resolution utility of primer sets for specific taxa. Secondly, the performance of the primers was evaluated *in vitro* using tissue derived DNA extracts with varying levels of degradation. Finally, eDNA samples, obtained from tanks of known species composition at the Aquarium of Genoa (Italy) as a proxy for ‘real world’ environmental samples, were used to assess the metabarcoding performance of the primers.

### *In silico* primer evaluation

An *in silico* approach was used to assess the universality of the newly designed primers against all standard nucleotide sequences in the NCBI-Genbank data repository (accessed April 2019) for three taxonomic groups: i) vertebrates (excluding Cetaceans), ii) Cetaceans only, and iii) Invertebrates. Performance of the newly designed primers were compared against the commonly used 12S-V5 vertebrate metabarcoding primers (Riaz et al. 2011) as a benchmark.

The ecopcr script was used to simulate an *in silico* PCR and extract the amplifiable barcoding regions for each primer pair while allowing for a maximum of 3 base-pair (bp) mismatches between the primers and template DNA. Barcode regions shorter than 50 bp and longer than 400 bp were not considered. Subsequently, the obigrep command was used to extract sequences that were reliably assigned to a species level taxonomy. Ambiguous species level identifications (i.e. sequences with ‘sp.’, ‘aff.’, etc. in the definition of the sequence) and nuclear pseudogenes were excluded from the final sequence database. The obiuniq command was then used to remove duplicate records for each species. Sequences were classified according to their higher taxonomy (i.e. Vertebrates [excluding Cetacean species], Cetaceans and Invertebrates) before summarising the data into a tabular format for further analyses using R version 3.5.2 (R Development Core Team 2010). Finally, the taxonomic resolution of the different primers was assessed by splitting the data into their higher taxonomy and running the ecotaxspecificity script with three different thresholds for barcode similarity (i.e. sequences were considered different if they have 1, 3 or 5 bp differences). The data obtained from the *in silico* analyses were imported into R and the packages tidyverse (Wickham 2016) and gridExtra (Auguie 2016) were used to construct summary figures to evaluate the taxonomic coverage, the specificity and taxonomic resolution for each primer pair.

We downloaded 680 GenBank complete marine mammal mitogenome sequences from GenBank, and evaluated levels of polymorphism within Families, Genera and species at the three proposed loci. Complete 12S and 16S genes were extracted from the retrieved sequences and aligned for each type of taxonomic comparison and the number of variable sites recorded within the three loci amplicons were reported.

### *In vitro* primer evaluation

Tissue derived DNA extracts were used to assess the performance of the newly designed primer pairs *in vitro* and optimise amplification conditions. DNA extracts of diverse marine vertebrate groups (cetaceans, pinnipeds, sea turtles and fish) were used as templates for PCR amplification (see Table 4). The 13 DNA templates were purposely selected to have different levels of degradation, being extracted with different techniques and spanning 1-31 years since extraction, in order to evaluate the ability of the primer to amplify low-quality DNA. High quality DNA extracts were obtained from fresh samples (i.e. muscle, skin or blood) using the Qiagen DNeasy Blood & Tissue extraction kit and following the manufacturers protocols. Low quality DNA extracts consisted of phenol-chloroform extracted DNA which was over 25 years old or DNA extracts obtained from boiling tissue samples in a buffer solution (Valsecchi 1998). For each primer pair and each of the DNA extracts a (single, duplicate, triplicate) PCR reaction was performed consisting of 25 U/μl of GoTaq G2 Flexi DNA polymerase (Promega); 1X Green GoTaq Flexi Reaction Buffer (Promega), 1.25-1.87 mM MgCl_2_ (Promega); 0.2 mM dNTPs (Promega), 0.25-0.75 μM of each primer and dH_2_O to reach a final volume of 20 μl. Thermal cycling conditions followed a touch-down PCR protocol with annealing temperatures depending on primer pairs: 10/10/18 cycles at 54/55/56 °C for MarVer1; 10/10/18 cycles at 51/52/53 °C for MarVer2; 8/10/10/10 cycles at 54/55/56/57 °C for MarVer3 and finally 10/10/18 cycles at 59/60/61 °C for Ceto2. After an initial denaturation step of 4 min at 94 °C, each of the 38 cycles consisted of a 30 s at 95 °C, 30 s at the primer specific annealing temperatures as described above, followed by 40 s at 72 °C. The final extension consisted of 5 s at 72 °C. To confirm the amplification of the desired amplicon, PCR products were visually assessed using gel-electrophoresis and Sanger sequenced (GENEWIZ, UK; data not shown).

### Evaluation of primer performance with environmental samples

Environmental DNA samples derived from water collected from 6 tanks of the Aquarium of Genoa, Italy in June 2018, were also employed to further validate the performance of the four primer sets. The tanks contained from 1 to 14 known vertebrate species and were named after their main hosted species or typology: 1) “Manatee”, 2) “Dolphin”, 3) “Shark”, 4) “Seal”, 5) “Penguin” and 6) “Rocky shore” - a multispecies tank hosting fish and invertebrates typical of Mediterranean rocky shores. Two tanks (Dolphin and Seal) were single species, while the Shark and the Rocky shore tanks included a combination of cartilaginous and bony fish.

For each tank, a total of 13.5 litres of water was collected from the water surface using a sterile graduated 2000ml glass cylinder, while wearing sterile gloves, and were stored within a 15-litre sterile container in order to homogenise the water sample, and avoid stochastic variability due to sampling of small volumes. For each tank, 3x 1.5 litre and 3x 3 litre replicates (total 6 replicates per tank) were then aliquoted from the larger sample. To capture environmental DNA, immediately after aliquoting, each of the 6 replicates was then filtered using individual 0.45 μm pore size nitrocellulose filters, using a BioSart^®^ 100 filtration system (Sartorius). After filtering, membranes were placed on ice for transport to the University of Milano-Bicocca, and subsequently stored at −20°C. Two weeks later environmental DNA was extracted from the filter membranes using a DNeasy PowerSoil Kit^®^ (Qiagen), following the manufacturer’s protocol.

For each of the four primer sets developed in this study, PCR performance with the eDNA extractions, was initially evaluated using the same PCR conditions as the *‘in vitro*’ validation. Subsequently two of the primer sets were selected for further use in a DNA metabarcoding analysis with high throughput sequencing using an Illumina MiSeq platform. For each selected primer set (metabarcoding locus), indexed forward and reverse sequencing primers were created comprising (5’ to 3’): an 8bp Illumina bar code tag - 4 random nucleotides - amplification primer sequence, and sourced from Sigma, UK. Eight forward primer indexes were combined with 12 reverse primer indexes, to allow pooling of amplicons from up to 96 uniquely identifiable samples within a single sequencing library (Taberlet et al. 2018). Trial amplifications with the Illumina barcode tagged primers suggested their yield and specificity was unchanged, and so the previously optimised PCR conditions were used to generate amplicons for MiSeq sequencing. For each locus, eDNA was amplified in triplicate in 40μl final PCR volume; 5μl of each replicate was used to check for successful amplification via agarose gel electrophoresis, and the remainder combined to yield a single pool for each sample.

Each sample amplicon pool was first assessed for fragment size distribution using an Agilent Tapestation, and cleaned with AMPure beads (Beckman Coulter), following the manufacturer’s protocol, to remove primer dimers. The cleaned samples were quantified with a Qubit fluorometer, and then for each metabarcoding locus, separate Illumina NEBNext Ultra DNA libraries were generated, with the pooled samples in equimolar ratios. Prior to sequencing each library was further assessed for fragment size distribution and DNA quantity by Agilent Tapestation and Qubit fluorometer. The library for locus MarVer1 (see results) was sequenced in a 150bp paired-end lane, and locus MarVer3 (see results) in a 250bp paired-end lane, using an Illumina MiSeq sequencer at the University of Leeds Genomics Facility, St James’s Hospital.

### Bioinformatics for environmental HTS data

Paired reads were first screened for the presence of the expected primer and index sequence combinations to exclude off-target amplicons. The reads were then combined to generate the insert sequence, and the sequence of the random nucleotide region noted, such that only one instance of an insert per sample with the sample random nucleotide finger print was saved to a sample specific file (i.e. to avoid PCR duplicates and chimeric sequences). The insert data was aggregated to create a counts matrix containing the occurrence of each unique sequence in each sample. The taxonomic origin of each insert was determined by Blasting their sequence against a local instance of the Genbank NT database (Nucleotide, [Internet]). The level of homology of insert to the hit sequence was noted, as was the species name of the hit sequence. The taxonomic hierarchy for each unique insert was generated by searching a local instance of the ITIS database (ITIS, [internet]) with the annotated Genbank species name. The count matrix and taxonomic hierarchy for all annotated unique sequences were then aggregated into values for equivalent Molecular Operational Taxonomic Units (MOTUs), by combining all inserts with a set homology (>=98%) to the Genbank hit at a specified taxonomic level (i.e. ‘Order’, ‘Family’, ‘Genus’ or ‘Species’), using bespoke software (available on request). Summaries and visualisations of read counts for different taxonomic levels were generated using the R package ‘Phyloseq’ (McMurdie and Holmes, 2013).

## RESULTS

### Description of primer sets

From the initial evaluation of aligned marine vertebrate mitochondrial sequences, four novel primer sets were identified, three within the 12S and one in the 16S genes (Table 1). The three ‘MarVer’ primer combinations (2 in 12S and 1 in 16S) were developed to amplify the target regions in any of the 71 taxa selected as representatives of marine vertebrates. To allow for variable sites between different vertebrate groups within the primer sequence, single degenerate bases were introduced (Table 1) for 4 out of the 6 MarVer primers. The fourth primer set ‘Ceto2’, was designed to preferentially amplify Cetacean DNA by minimizing base-pair mismatches for cetacean species while maximizing base-pair mismatches for other vertebrate groups. Online Resource 2 (Supplementary Material) shows variability at the priming sites across the eight marine vertebrate categories. Amplicon variability across the 71 taxa (plus the 4 human sequences) recorded in the regions targeted by the proposed markers are shown in Online Resource 3 (Supplementary Material).

**Table 1.**
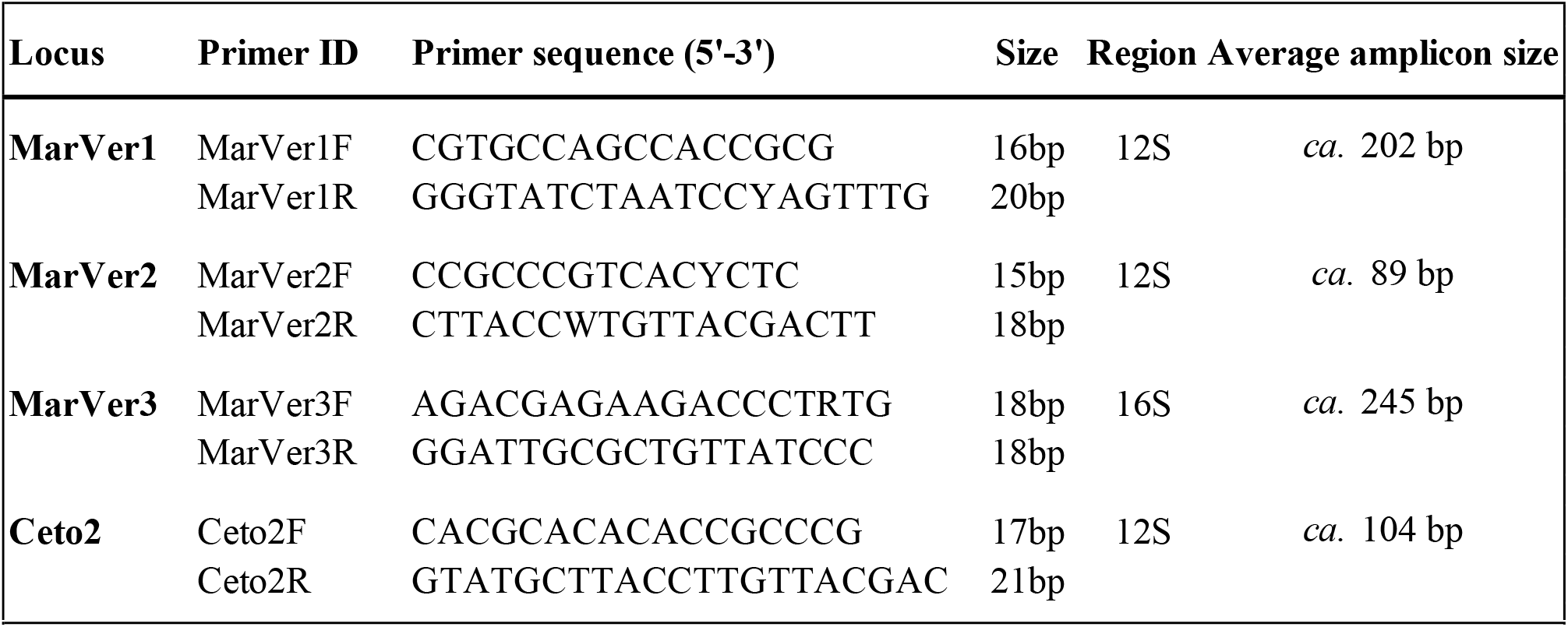
Sequences of the four described primer sets

#### MarVer1

MarVer1 primer set (abbr. MV1) targets a hypervariable region of the 12S gene, amplifying a fragment of about 199-212bp (Table 2). It partially overlaps with loci Tele02 (Taberlet et al. 2018), Tele03 (as named by Taberlet et al. 2018, corresponding to MiFish-U in Miya et al. 2015) and Elas01 (as named by Taberlet et al. 2018, corresponding to MiFish-E in Miya et al. 2015), targeting bony and cartilaginous fishes (Figure 1). The forward primer MarVer1F differs from the forward primers of previously described loci, in that by shifting 5-12bp at the 5’ end, it skips variable sites distinguishing bony from cartilaginous fishes, while gaining, at the 3’ end, bases which are conserved across all surveyed marine vertebrates.

**Table 2.** Characteristics of the amplicons produced by each of the primer sets for the 71 marine vertebrate species used for primers’ design. The overall amplicon size is the mean value of the amplicon sizes recorded in the eight marine vertebrate groups analysed. Ceto2 locus is not included in this table as its characteristics are the same as MarVer2, with the only exception of having the total amplicon size increased of 15bp compared with MarVer2 due to primers positioning. The grey shading indicates, for each vertebrate group, rank of nucleotide p-distance values across the three loci (the highest shown in darker grey)

| **MarVer01** | **CODO** | **CMYS** | **PINN** | **SIRE** | **STUR** | **SBIR** | **TELE** | **ELAS** | **Overall** |
| --- | --- | --- | --- | --- | --- | --- | --- | --- | --- |
| approximate amplicon size (bp) | 199 | 199 | 197 | 199 | 211 | 209 | 197 | 212 | **ca. 202** |
| n variable sites | 77 | 31 | 33 | 18 | 36 | 45 | 80 | 40 | **171** |
| resolved over tested species | 36/36 | 12/12 | 3/3 | 2/2 | 6/6 | 2/2 | 7/7 | 3/3 | **71/71** |
| percentage of variable sites (%) | 38.7 | 15.6 | 16.8 | 9.0 | 17.1 | 21.5 | 40.6 | 18.9 | **84.7** |
| nucleotide p-distance | 0.084 | 0.042 | 0.097 | 0.084 | 0.059 | 0.157 | 0.176 | 0.090 | **0.196** |
| **MarVer02** | **CODO** | **CMYS** | **PINN** | **SIRE** | **STUR** | **SBIR** | **TELE** | **ELAS** | **Overall** |
| approximate amplicon size (bp) | 96 | 96 | 89 | 90 | 87 | 87 | 85 | 85 | **ca. 89** |
| n variable sites | 46 | 22 | 29 | 16 | 18 | 11 | 47 | 26 | **65** |
| resolved over tested species | 30/36 | 12/12 | 3/3 | 2/2 | 6/6 | 2/2 | 7/7 | 3/3 | **65/71** |
| percentage of variable sites (%) | 47.9 | 22.9 | 32.6 | 17.8 | 20.7 | 12.6 | 55.3 | 30.6 | **73.0** |
| nucleotide p-distance | 0.056 | 0.028 | 0.146 | 0.125 | 0.075 | 0.047 | 0.175 | 0.109 | **0.154** |
| **MarVer03** | **CODO** | **CMYS** | **PINN** | **SIRE** | **STUR** | **SBIR** | **TELE** | **ELAS** | **Overall** |
| approximate amplicon size (bp) | 233 | 232 | 240 | 235 | 240 | 246 | 274 | 265 | **ca. 245** |
| n variable sites | 79 | 41 | 53 | 36 | 50 | 31 | 142 | 37 | **251** |
| resolved over tested species | 34/36 | 12/12 | 3/3 | 2/2 | 6/6 | 2/2 | 7/7 | 3/3 | **69/71** |
| percentage of variable sites (%) | 33.9 | 17.7 | 22.1 | 15.3 | 20.8 | 12.6 | 51.8 | 14.0 | **102.4** |
| nucleotide p-distance | 0.052 | 0.040 | 0.098 | 0.081 | 0.054 | 0.058 | 0.151 | 0.046 | **0.167** |

**Fig. 1.**
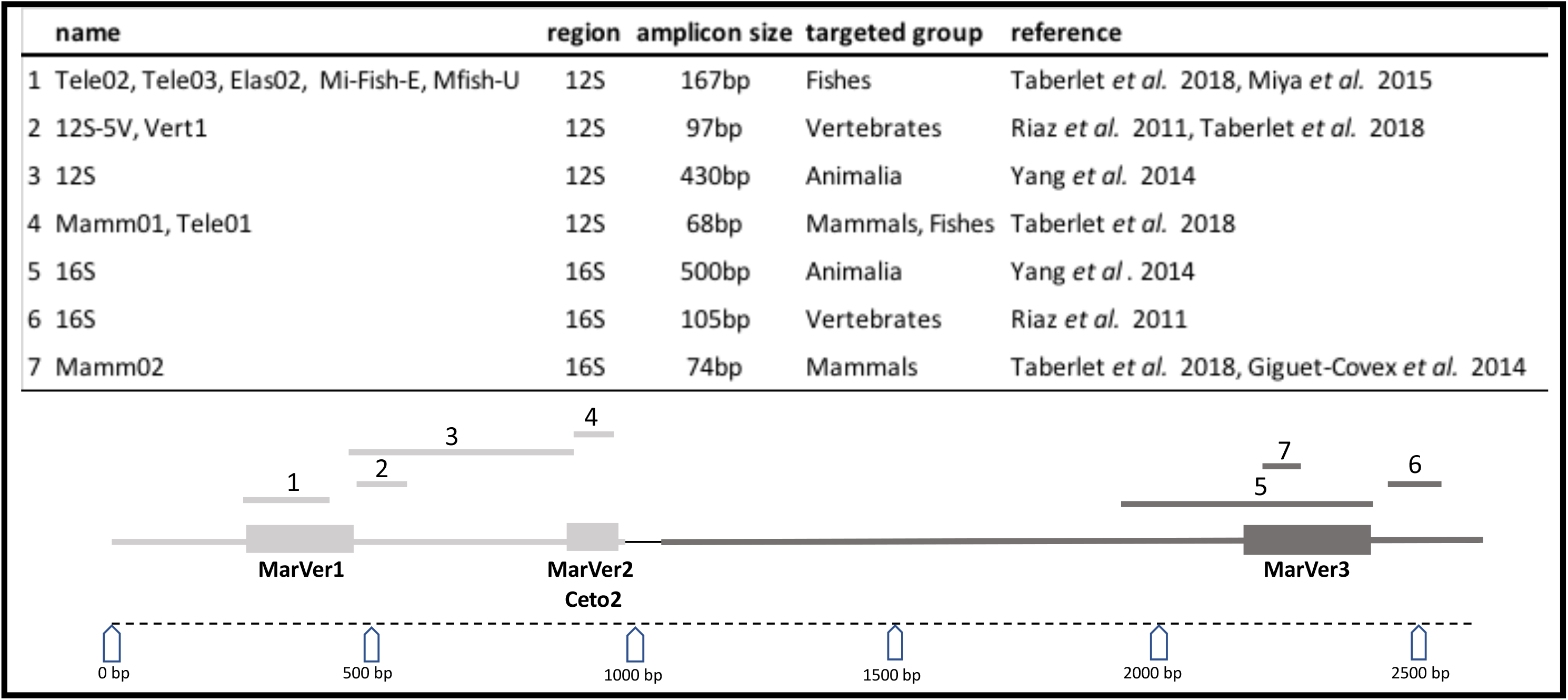
Map of the regions amplified by the newly presented primer sets within the 12S (light grey) and 16S (dark grey) genes. The positions of some of most commonly used barcode markers used for detecting vertebrate groups are shown for comparison. The size marker at the bottom refers to the 12S and 16S mtDNA fragment from position 72 to position 2690 in the stripe dolphin (*Stenella coeruleoalba*) complete mitogenome (GenBank accession number NC_012053)

#### MarVer2

MarVer2 primer set (abbr. MV2) targets a variable region of the 12S gene, amplifying a smaller fragment of 85-96bp in the tested vertebrate taxa (Table 2). It is a modification of primer sets Tele01 (Valentini et al. 2016), targeting bony fishes, and Mamm01 (Taberlet et al. 2018), targeting mammals (Figure 1). The MarVer2 reverse primer partially overlaps with AcMDB01R primer designed by Bylemans et al. 2018 for freshwater fish Actinopterygii species. Both forward and reverse MarVer2 primers were designed with a degenerate base to allow amplification in all the 71 tested vertebrate taxa (Table 2). Sequence variation within MarVer2 amplicon did not resolve all 71 marine vertebrate taxa in the original alignment, with sequences shared between two or more Odontocete species within the Delphinidae family; with *Sousa chinensis, Tursiops truncatus, Tursiops aduncus, Stenella coeruleoalba, Delphinus delphis* and *Delphinus capensis* sharing one amplicon sequence, and *Peponocephala electra* and *Feresa attenuaae* sharing another.

#### MarVer3

MarVer3 (abbr. MV3) amplifies a variable region of the 16S gene, producing amplicons ranging in size from 232 to 274bp in the 71 marine vertebrate taxa tested (Table 2). MarVer3 is partially covered by locus Mamm02 (Taberlet et al. 2018, see Figure 1), but targets a fragment twice as long: for example, in Odontocetes MarVer3 amplifies a 233bp fragment versus a 115bp amplicon for Mamm02. The reverse primer (MarVer3R) was the only one of the presented oligonucleotides to be truly “universal”, as it was found to be fully conserved across all tested marine vertebrates (Table 2). The MarVer3 amplicon sequence resolved 69 of the 71 tested marine vertebrate species. The unresolved species also fall into the Delphinidae family: *Sousa chinensis* and *Tursiops truncatus* sharing one amplicon sequence, and *Tursiops aduncus* and *Delphinus capensis* sharing another.

#### Ceto2

Ceto2 primers were designed to complement the MarVer2 locus, amplifying the same 12S region, but with nucleotides specific to Cetaceans (and excluding humans) in both forward and reverse primers, and containing no degenerate bases (Table 1). Ceto2 amplicon sizes were 15bp longer than MarVer2, due to shifting of the priming sites. The Ceto2 primer set is suggested as not suitable for use with marine mammals other than Cetaceans due to primer site mismatches. The priming sites present one variable site in some Pinnipeds with the New Zealand fur seal (*Arctocephalus forsteri*) exhibiting a variant at the 3’ end of the forward primer, while both the walrus (*Odobenus rosmarus*) and the harp seal (*Phoca groenlandica*) presented a variant towards the 5’ end of the Ceto2R primer.

### Primer evaluation and validation

#### *In silico* primer evaluation

For the different primer pairs and higher taxonomic groups, the total number of unique taxa for which the *in silico* amplification recovered target sequences is given in Figure 2. The results show that the MarVer3 primer pair amplified DNA from the most vertebrate and cetacean taxa. However, this primer pair also successfully amplifies the DNA of invertebrate species thus indicating that it has a low overall specificity to the intended taxonomic targets (Figure 2). The three other primer pairs all amplified DNA from fewer target taxa then the commonly used 12S-V5 primers.

**Fig. 2.**
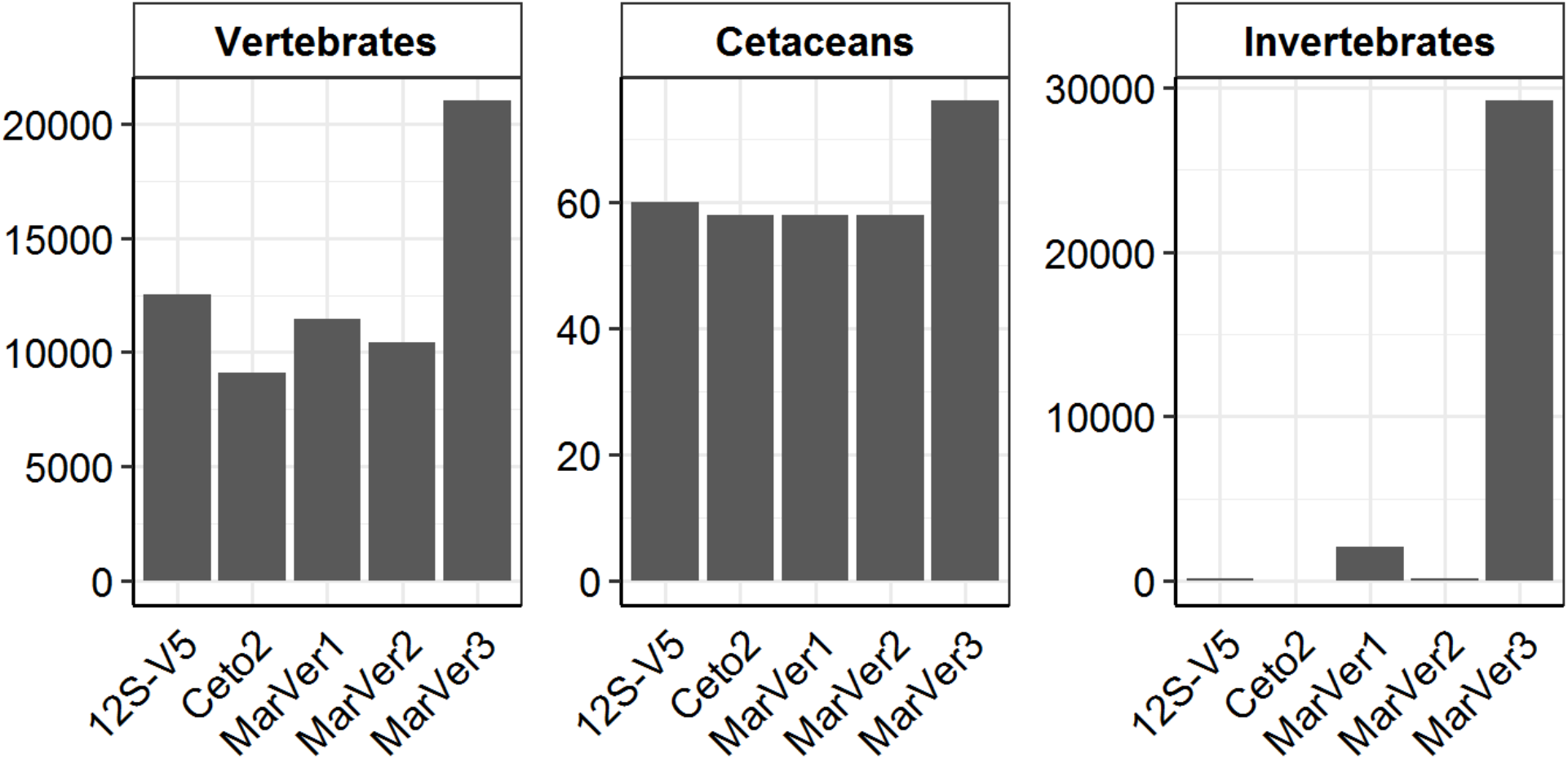
The total number of unique taxa (y-axis) recovered by the different primer pairs (x-axis). The results are shown for all three higher taxonomic groups (i.e. Vertebrates -excluding cetacean species-, Cetaceans and Invertebrates) considered during the analyses (horizontal panels)

The proportion of sequences amplified for each higher taxonomic group is a function of the bp-mismatches between the primers and template DNA is shown in Figure 3. The results of the commonly used 12S-V5 primer pair show that a high number of vertebrate sequences have very few bp-mismatches while the recovered non-vertebrate sequences generally have ≥ 5 bp-mismatches at both primer binding regions (Figure 3). A similar pattern can be observed for the MarVer2 primer pair although the bp-mismatches in the primer binding regions for non-vertebrate sequences are slightly lower (i.e. ≥ 4 bp-mismatches between both primer regions). The results of both the MarVer1 and MarVer3 primers shows that even with a low number of bp-mismatches (i.e. ≥ 2 mismatches for both the forward and reverse primers) a significant proportion of the amplified sequences belong to non-vertebrate taxa. Finally, the results of the Ceto2 primers show that this primer pair will preferentially amplify Cetacean species as the bp-mismatches between Cetacean sequences are lower compared to the number of bp-mismatches observed in the primer binding regions for most vertebrate sequences (i.e. ≤ 2 bp-mismatches for the Cetacean sequences versus ≥ 3 bp-mismatches for most vertebrate sequences).

**Fig. 3.**
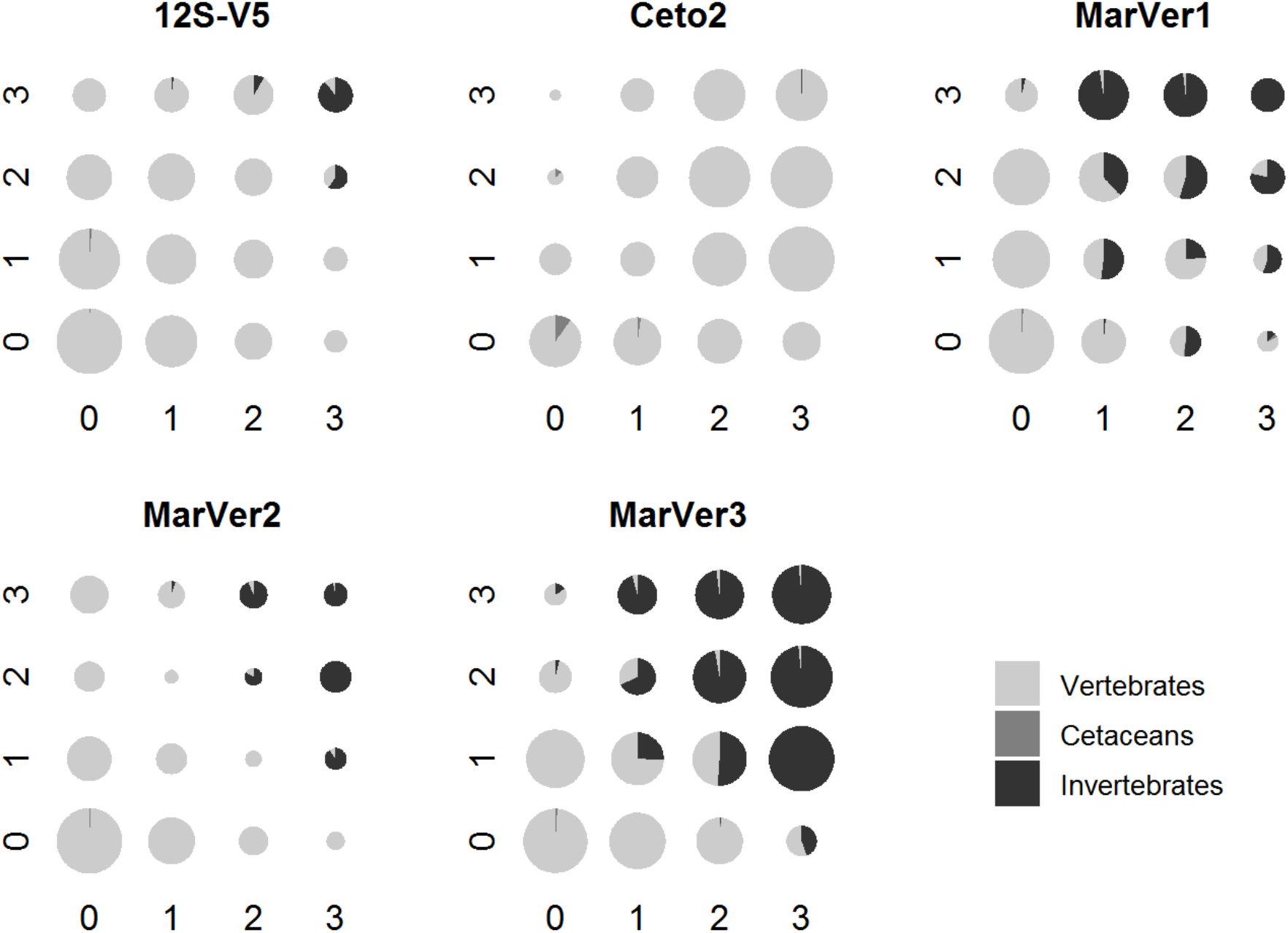
The proportion of sequences amplified for each higher taxonomic group as a function of the bp-mismatches between the template DNA and the forward (y-axis) and reverse (x-axis) primers. The size of the pie-charts is proportional (on a log scale) to the total number of sequences recovered for a given number of bp-mismatches between the forward and reverse primer

The taxonomic resolution power of the different primer pairs was evaluated using both the Cetacean and Vertebrate sequences and the results are summarised in Figure 4. Overall the commonly used 12S-V5 metabarcoding primers have a lower resolution capacity compared to our newly designed primers (for both the Cetacean and Vertebrate taxa), with our primers assigning 20-25 % more sequences to the correct species level taxonomy (Figure 4). For the newly designed primers, differences are observed in their taxonomic assignment power, with MarVer1 and MarVer3 generally assigning a higher percentage of the sequences to the correct family, genus and species level taxonomy (Figure 4).

**Fig. 4.**
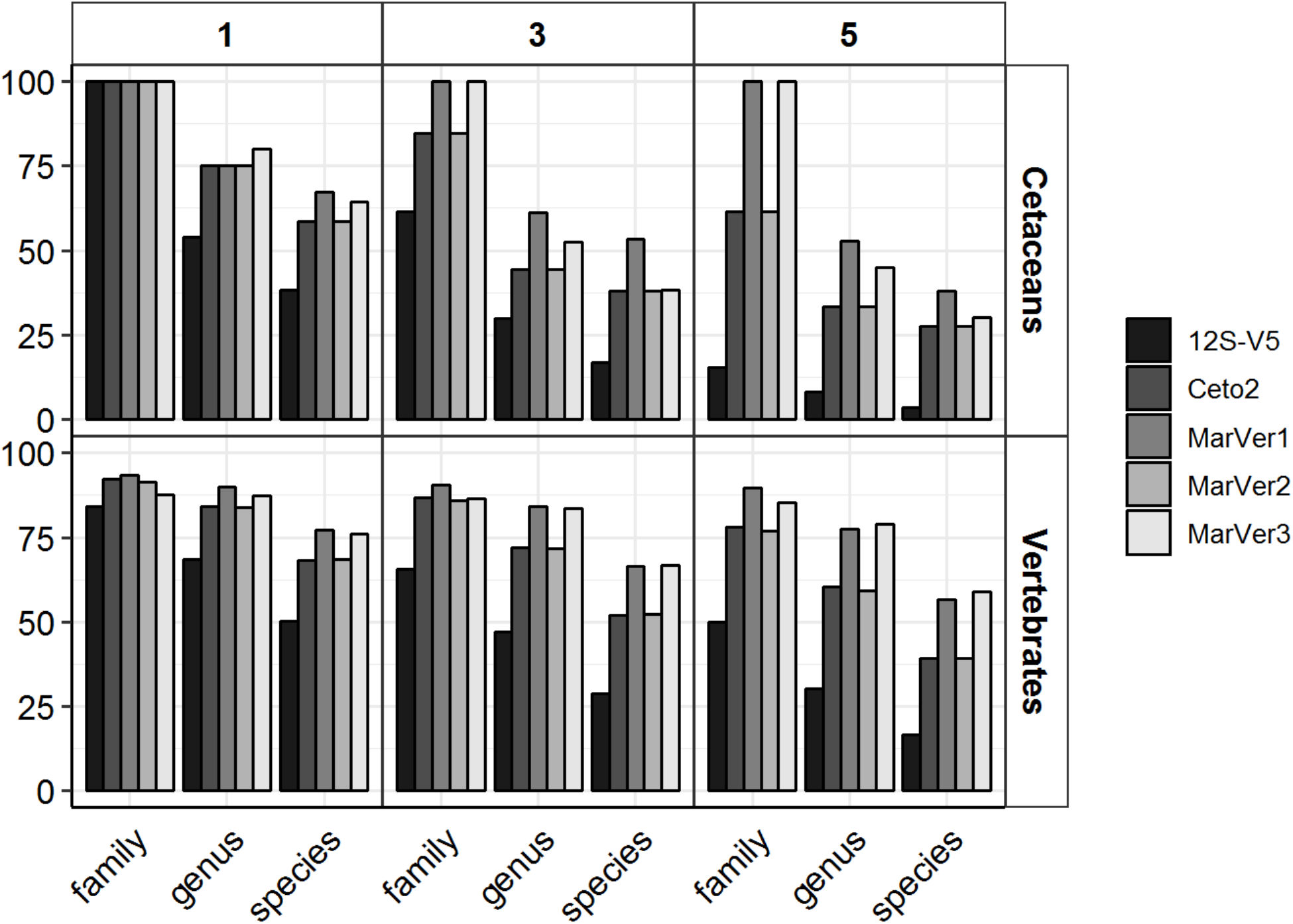
The percentage of sequences correctly identified (y-axis) to the family, genus and species level taxonomy (x-axis) for the different primer pairs. Results are shown for all sequences belonging to the Cetaceans and Vertebrates (vertical panels) and using different threshold values for barcode similarity (i.e. sequences were considered different if they have 1, 3 or 5 bp differences) (horizontal panels)

#### Primer set resolution for Marine Mammal taxonomic detection

Table 3 shows the results of the comparison performed on 680 GenBank complete marine mammal mitogenome sequences (GenBank accession numbers shown in Online Resource 4), in order to evaluate levels of polymorphism within Families, Genera and species.

**Table 3.**
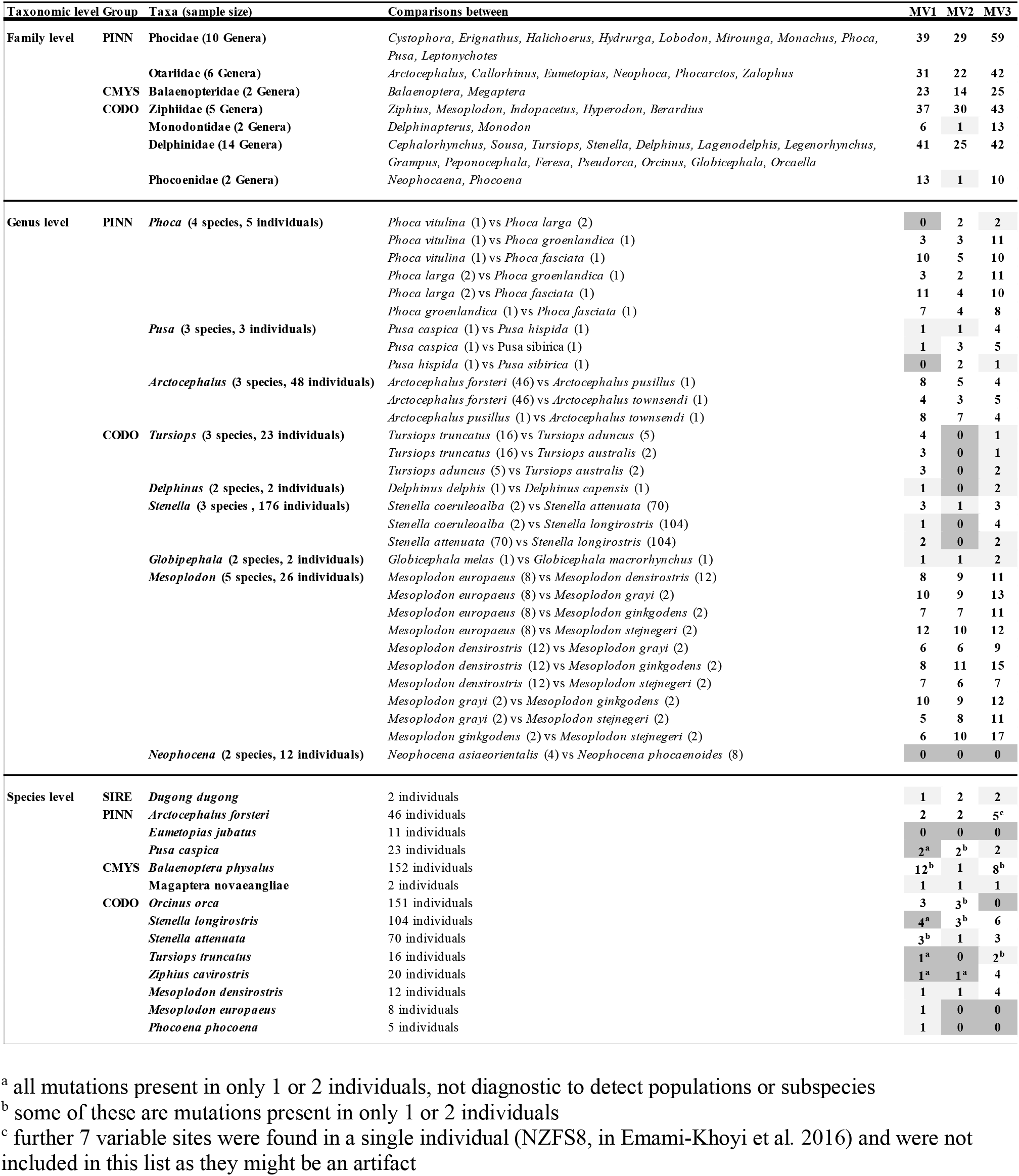
Levels of polymorphism found at the three proposed loci in a series of marine mammals Families, Genera and species for which complete mitogenomic data were available from GenBank (accession numbers listed in Appendix E, with the exception of 22 *Pusa capsica* entries for which data are unpublished). Numbers in the three columns on the right indicate the number of variable sites detected in the sequence targeted by the three primers sets in each given comparison: clear boxes indicate polymorphism thus usefulness of the primer set to resolve taxa case, while grey boxes show lack of diagnostic polymorphism and light grey boxes indicate lack of resolution at 98% homology threshold for MOTUs assignment

##### Family level

All three targeted regions contained high genetic variability within the seven analysed marine mammals Families (Pinnipeds (2), Mysticetes (1) and Odontocetes (4)). The DNA fragment amplified by MarVer3 primer set (16S region) proved to be the most diverse, highlighting 59 variable sites within the Phocoidea, and over 40 in the Otariidae, Ziphiidae and Delphinidae Families. The least variable locus of the three was MarVer2 (thus also Ceto2). Nevertheless, this marker had high discriminatory power for genera within Families, with the exception of the Monodontidae and Phocoenoidae families. Here both families contain only 2 genera, which in both cases only differ by one basepair, and therefore may not be automatically resolved as distinct MOTUs if a threshold of 98% homology is used in the annotation pipeline (Table 3).

##### Genus level

Nine Genera, each including 2 to 5 species, were assessed for within-genus variability, including pairwise con-generic species comparisons (Table 3). MarVer3 consistently revealed the highest levels of polymorphism (23 of the 31 inter-generic pairwise comparisons show differences of at least 1bp), while MarVer2 was least variable, showing no variation in 7 of the 31 pairwise comparisons (grey boxes in Table 3). However, MarVer2 showed good power to resolve taxon identity in the Genus *Mesoplodon* (Ziphiidae family). MarVer1 was the second most variable locus at this level, with the highest number of variable sites in 10 out of the 31 congeneric-species pairwise comparisons. Within all three *Tursiops* spp. pairwise comparisons MarVer1 was the locus showing the highest variability.

##### Species level

Genetic variability was investigated also below the nominal species level. This could be tested only on the few species for which large enough sample sizes were available to evaluate intraspecific variation. We assessed 14 marine mammal species for which mitogenomic data were available for a number of individuals ranging from 2 (*Megaptera novaeangliae* and *Dugong dugong*) to 152 (*Balaenoptera physalus*) individuals (Table 3). The 14 species were representative of seven marine mammal Families: Dugongidae (1 species), Otariidae (2 species), Phocidae (1 species), Balaenopteridae (2 species), Delphinidae (4 species), Ziphiidae (3 species) and Phocoenidae (1 species). All sequences were retrieved from GenBank (see Online Resource 4), with the exception of 22 unpublished *Pusa caspica* sequences (provided by SG). In only one of the 14 species (*Eumetopias jubatus*, fam. Otariidae) were none of the three loci polymorphic (11 individuals compared). Four of 14 species had only one informative locus: MarVer1 (12S) in two instances (*Mesoplodon europaeus* and *Phocoena phocoena*), and MarVer3 (16S) in two other cases (*Tursiops truncatus* and *Ziphius cavirostris*) (Table 3). In the Caspian seal (*Pusa capsica*), the killer whale (*Orcinus orca*) and the spinner dolphin (*Stenella longirostris*), two of three loci showed polymorphism. In the remaining 6 species (*Dugong dugong, Arctocephalus forsteri, Balaenoptera physalus, Megaptera novaeangliae, Stenella attenuata, Mesoplodon densirostris*) all 3 loci presented polymorphisms: in some instances MarVer1 was found to be the most informative locus (e.g. 12 variable sites in *Balaenoptera physalus*), in others MarVer3 (e.g. 6 variable sites in *Stenella longirostris*).

#### *In vitro* primer evaluation

PCR amplicons of the expected size were generated by the four primer sets with most of the 13 DNA extracts (Table 4). The oldest DNA extract (31yo, humpback whale, *Megaptera novaeangliae*) was the worst performing, yielding only very faint bands for MarVer1, MarVer3 and Ceto2, and failing to produce a band for locus MarVer2. All sea turtle extracts produced one strong artefact band (ca 161bp) and a fainter band in the correct size range (ca 107bp) place of the correct sized fragment for locus Ceto2. Bony fish DNA extracts also produced a band compatible in size with the Ceto2 amplicon.

**Table 4.**
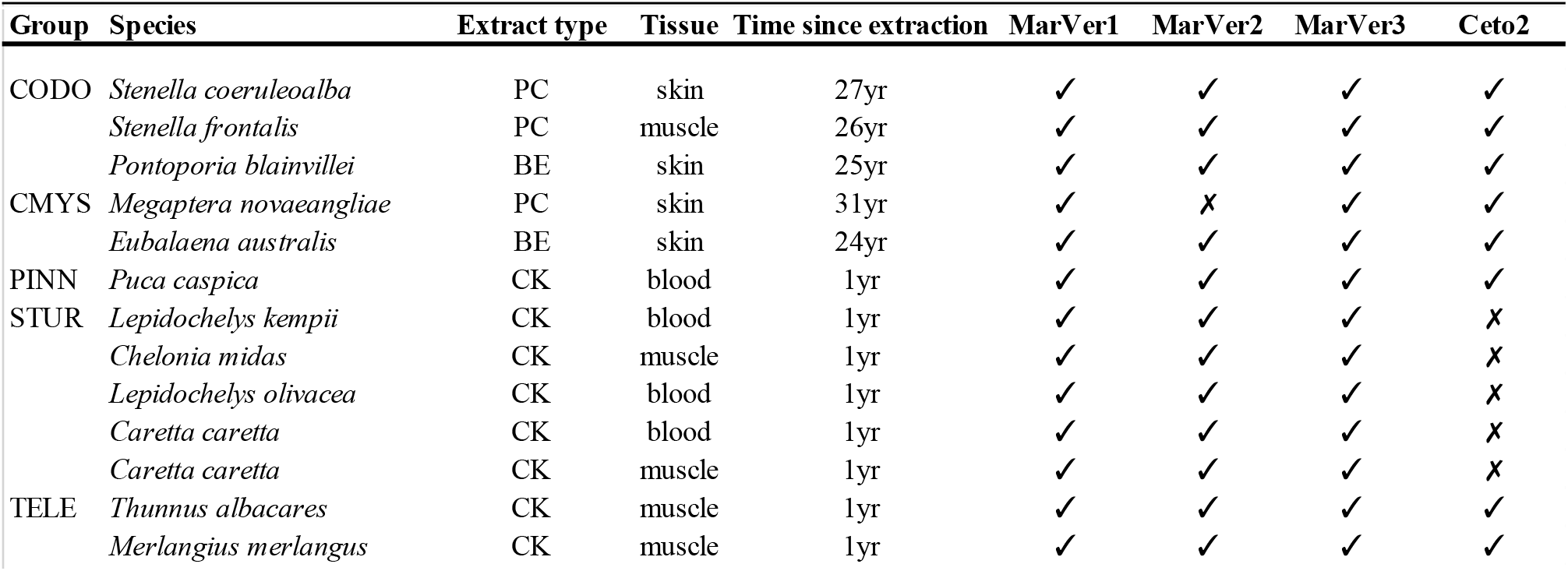
List of tissue DNA extracts used for wet lab primer tests. “PC” indicates phenol chloroform extracts, “CK” indicates commercial kit extracts, while “BE” refers to boil extracts as described in Valsecchi 1998. The signs “✓” and “✗” specify successful amplification and PCR failure (no band or aspecific bands) respectively

#### Application to environmental samples

Amplicons of the expected size were obtained from all 36 water samples collected at the Genoa Aquarium with the four primer sets. Locus Ceto2 amplified a faint band in the expected size range from the shark, penguin and seal tanks, in addition to the bottlenose dolphin tank, suggesting either low level amplification from non-cetacean taxa or potential transfer of dolphin eDNA between tanks (see below).

Loci MarVer1 and MarVer3 were selected for further HTS metabarcoding evaluation on the basis of the bioinformatic prediction that they would be the most informative for Mediterranean marine megafauna species (see below). For these loci after sequence quality filtering and demultiplexing, annotated read counts per sample ranged from 682 to 52,478 for MarVer1 and 1,025 to 43,003 for MarVer3, with combined reads per Tank of 10,750 to 158,950 for MarVer1, and 1,3251 to 232,015 for MarVer3 (see Table 5, Figure 5, Online Resource 5).

**Table 5.** Species composition of the 6 tanks of the Aquarium of Genoa from which water samples were collected for extracting eDNA. The last four columns on the right show NGS outcomes for primer sets MarVer1 and MarVer3: “GB” columns indicate whether reference sequences are present in GenBank for that specific species and locus; “■” indicate successful species detection (half-full and empty square indicates a number of reads 0.001>nr>0.005 or nr<0,001 respectively); Δ denotes that a congenereric was identified in place of the resident vertebrate species, “?” indicates possible detection of the resident species at a higher taxonomic group (this case together with those instances for which reference sequences were available but the species remained undetected are discussed in the text)

| Tank | Name | Hosted species (n. individuals) | Common name | Vertebrate group | MarVer1 |  | MarVer3 |  |
| --- | --- | --- | --- | --- | --- | --- | --- | --- |
|  |  |  |  |  | GB | detection | GB | detection |
| 1 | MANATEE | *Trichechus manatus* (4) | Matatee | SIRE | yes | ■ | yes | ■ |
|  |  | *Piaractus brachypomus* (3) | Pirapitinga | TELE | yes | ■ | yes | ■ |
|  |  | *Pterygoplichthys gibbiceps* (2) | Armored catfish | TELE | no | X | yes | X |
| 2 | DOLPHIN | *Tursiops truncatus* (10) | Bottlenose dolphin | CODO | yes | ■ | yes | ■ |
| 3 | SHARK | *Ginglymostoma cirratum* (3) | Nurse shark | ELAS | yes | ■ | yes | ■ |
|  |  | *Stegostoma fasciatum* (1) | Zebra shark | ELAS | yes | ■ | yes | ■ |
|  |  | *Carcharhinus plumbeus* (4) | Grey shark | ELAS | yes | ■ | yes | ■ |
|  |  | *Dasiatis americana* (1) | Southern stingray | ELAS | no | X | no | X |
|  |  | *Pristis zijsron* (2) | Longcomb sawfish | ELAS | yes | X | yes | X |
|  |  | *Teaniura grabata* (2) | Round stingray | ELAS | no | X | yes | X |
|  |  | *Epinephelus itajara* (1) | Atlantic goliath grouper | TELE | no | △ | yes | ☑ |
|  |  | *Epinephelus costae* (1) | Goldblotch grouper | TELE | no | △ | yes | ■ |
|  |  | *Epinephelus marginatus* (2) | Dusky grouper | TELE | no | △ | yes | ■ |
|  |  | *Seriola dumerili* (9) | Greater amberjack | TELE | yes | ■ | yes | ☑ |
|  |  | *Dicentrarchus labrax* (7) | European seabass | TELE | yes | ■ | yes | ■ |
|  |  | *Diplodus sargus* (4) | White seabream | TELE | no | X | yes | ■ |
|  |  | *Diplodus cervinus* (1) | Zebra sea bream | TELE | no | X | yes | △ |
|  |  | *Pagellus bogaraveo* (1) | Blackspot seabream | TELE | yes | X | yes | □ |
| 4 | PENGUIN | *Pygoscelis papua* (12) | Gentoo penguin | SBIR | yes | X | yes | ■ |
|  |  | *Spheniscus magellanicus* (24) | Magellanic penguin | SBIR | yes | △ | yes | ■ |
| 5 | SEAL | *Phoca vitulina* (6) | Harbour seal | PINN | yes | ■ | yes | ■ |
| 6 | ROCKY SHORE | *Diplodus vulgaris* (21) | Common two-banded seabream | TELE | no | X | yes | ■ |
|  |  | *Diplodus sargus* (3) | White seabream | TELE | no | X | yes | ■ |
|  |  | *Sciaena umbra* (3) | Brown meagre | TELE | no | ? | yes | ■ |
|  |  | *Sarpa salpa* (2) | Salema | TELE | no | X | yes | ■ |
|  |  | *Serranus scriba* (1) | Painted comber | TELE | no | X | yes | ☑ |
|  |  | *Scyliorhinus stellaris* (2) | Nursehound | ELAS | no | X | yes | ■ |
| Feed species |  |  |  |  |  |  |  |  |
| 2,3,4,5,6 |  | *Culpea harengus* | Atlantic herring | TELE | yes | ■ | yes | ■ |
| 2,3,4,5,6 |  | *Mallotus villosus* | Capelin | TELE | yes | ■ | yes | ■ |
| 3,6 |  | *Merluccius productus* | North Pacific hake | TELE | no | X | yes | ■ |
| 6 |  | *Scomber scombrus* | Atlantic mackerel | TELE | yes | ☑ | yes | ■ |
| 2 |  |  | Squid |  | yes |  | yes |  |

**Fig. 5.**
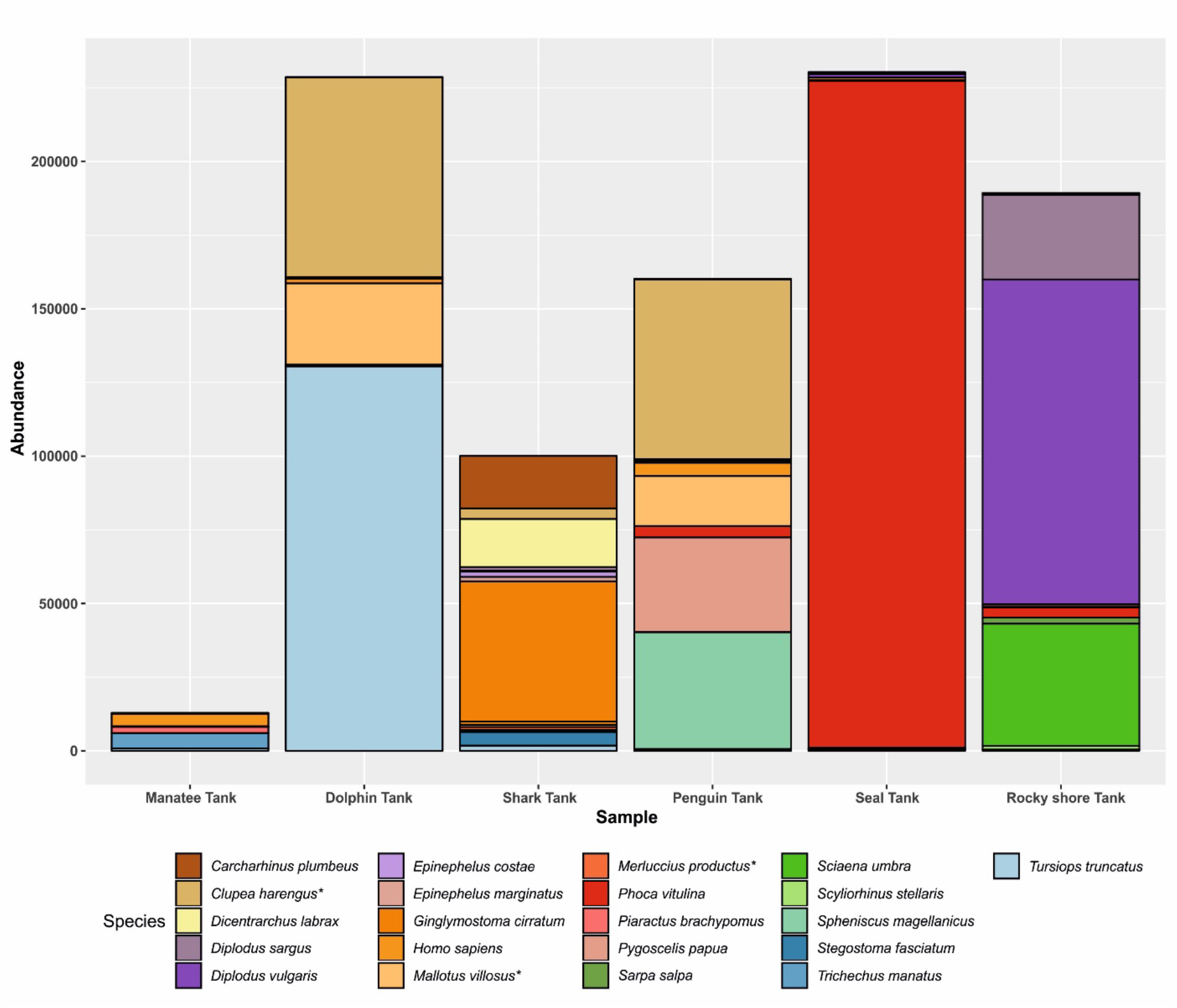
Taxon ‘abundance’ (read counts) for amplicons generated with the MarVer3 primer set for environmental DNA extracted from water samples collected from 6 tanks at the Genoa Aquarium. Read counts are combined across the 6 replicate samples assayed for each tank, and seqeunce demultiplexing and amplicon annotation against Genbank references was carried out using a threshold of 98% sequence similarity. Taxa presented in the figure were filtered to exclude those with read counts less than 0.005*median read count across tanks. Taxa names with *, are feed species, all others are resident

The percentage of resident vertebrate species with amplicons annotated to the species level within each tank ranged between 0% (Tank 6, Rocky shore) to 100% (Tank 2, Dolphin and Tank 5, Seal), mean 50.4% for MarVer1; and between 66.6% (Tank 1, Manatee) to 100% (Tanks 4, 5, 6), mean 89.7%, for MarVer3 (see Online Resource 5). Overall, for MarVer1 amplicons for 9 out of 27 resident taxa were annotated to the species level, and for 22 resident species for MarVer3, (see Table 5, Figure 5, Online Resource 5).

Amplicons for the aquarium’s four frequently used vertebrate feed species (*Culpea harengus, Mallotous villosus, Merluccius productus*, and *Scomber scombrus*), were detected in tanks 2 to 6 (in the Manatee tank, feed consists of lettuce) using both loci, with the exception of MarVer1 failing to detect *Merluccius productus*. Squid (unspecified species), is also supplied to the Dolphin Tank, but no cephalopod, amplicons were recovered.

Amplicons for resident species were also detected in tanks other than their host tanks at low levels (e.g. with MarVer3, dolphin and seal amplicons were detected in the manatee and shark, and penguin and rocky shore tanks respectively), suggesting possible transfer of eDNA between tanks in the aquarium, e.g. via the equipment used by staff members. Similarly, human amplicons were detected in all tanks, consistent with the practice of aquarium staff entering the water for maintenance.

Both MarVer1 and MarVer3 identified amplicons (partially overlapping between loci and tanks) that were not directly attributable to resident species or food sources (category B in Online Resource 5). These comprised 6 recurrent species, two of which were previously (but no longer) used as feed (*Sardina pilchardus, Engraulis encrasicolus*), and four species present in the Ligurian Sea (e.g. *Auxis rochei, Auxis thazard, Belone belone, Coris julis*) from which the water used to fill the tanks is drawn, after being filtered and UV irradiated. All of these unexpected species detections were at low abundance, with read counts greater than 100 to a maximum of 947 in at least 1 tank (range 0.3% to 3.7% of total reads with MarVer3 and MarVer1 respectively). Very low abundance amplicons (<100 reads per tanks) for at least 20 other Mediterranean resident teleost fish species were also observed, but were not considered further as definitive detections.

Amplicons from invertebrate species (category C in Online Resource 5) were detected in two tanks by the MarVer3 locus at low abundances (read count <100), which would normally be discounted as a detection. These being, an Anthozoa species, *Seriatopora hystrix*, in the manatee tank, and a Sipunculid worm, of the family Phascolosomatidae, in the seal tank (Online Resource 5). Neither taxa were in the tank in which their traces were found. No invertebrate amplicons were recovered in the “rocky shore” tank which contains some Anthozoa (e.g. *Anemonia viridis*), some unidentified sponges growing spontaneously and hydrozoans, or from the shark tank where *Aiptasia* spp grows spontaneously. The other tanks (with the exception of dolphin and manatee) may also contain other spontaneously growing small invertebrates, such as copepods, amphipods and hydrozoans, but none were detected.

## DISCUSSION

### Comparison with previously described barcode primer sets

This study was conceived to identify cetacean specific barcode loci complementing the many already available for fish species (e.g. Miya et al. 2015, Sato et al. 2018), for use in eDNA biodiversity monitoring studies of Mediterranean marine vertebrates. However, in the primer design process, we realised that with minor adaptations, it was possible to cover the whole range of marine vertebrates in a single HTS run, potentially dramatically reducing costs for eDNA HTS biodiversity monitoring of pelagic vertebrates.

Most existing 12S/16S barcode primer sets (e.g. Taberlet et al. 2018, Miya et al. 2015, Bylemans et al. 2018) were designed for particular vertebrate groups and thus contained conserved sequence elements specific for their target taxonomic group. Most of these primer sets are partially overlapping, at least at one end, with the ones presented in this study although they are never identical (Figure 1). Our proposed primer sets were also found to be different from the universal 12S and 16S primers combinations described by Yang et al. (2014) - although MarVer3 sits within Yang et al.’s 16S target fragment - however, their amplicon sizes were too large (ca 430bp and ca 500bp for respectively the 12S and the 16S primer sets) to be easily employed in eDNA studies using current short-read HTS technology.

The 12S-V5 primer set (Riaz et al. 2011, renamed Vert01 by Taberlet et al. 2018), is the only one previously described as being specific for vertebrates. It is located adjacent to MarVer1 site (forward 12S-V5 primer partially overlaps with reverse MarVer1 sequence, see Figure 1). Within our alignments, the 12S-V5 site was not as variable as any of the three loci described in this paper. For example, looking at one of the most variable Families considered in this study, the Ziphiidae (Odontocetes), our three loci (MarVer 1, 2 and 3) recovered 37, 30 and 43 variable sites respectively (Table 3), while 12S-V5 highlighted only 13 within the same pool of sequences. Given that 12S-V5 and MarVer2 primer pairs generate similar sized amplicons (respectively 106bp and 96bp), MarVer2 detects more than twice (30) the number of variable sites found in the 12S-V5 target sequence (13). Such performance difference was also highlighted in the *in silico* simulation (Figure 4). Moreover, within both forward and reverse 12S-V5 primer sites, polymorphisms were present within the Ziphiidae Family, suggesting that the 12S-V5 primers may not be conserved across all vertebrate classes. However, 12S-V5 performed better than MarVer1, MarVer2 and Ceto2 based on the total number of unique taxa for which DNA was amplified in the *in silico* simulation (Figure 2). While the lower predicted number of taxa recovered by these primers may indicate a failure to amplify DNA from some species, the completeness of the reference database used will also influence the results given that all sequences records were considered and not only the full mitochondrial genomes. Within the partial 12S sequences deposited on GenBank the fragment including locus 12S-V5 might be over-represented, as most previous studies relied on this marker.

Besides being highly conserved among Vertebrates, all three MarVer primer-sets were shown to be potentially non-exclusive to Vertebrates when 2-4bp primer/template mismatches were allowed (Figure 3). With reduced specificity, these primer pairs could potentially amplify unwanted non-vertebrate taxa. Given the high number of vertebrate taxa recovered by the MarVer3 primers *in silico* (Figure 2), and the observation that the majority of the non-vertebrate taxa recovered have ≥ 3 bp-mismatches in the primer binding regions (Figure 3), this primer pair should still be valuable if stringent thermal cycling conditions are used during PCR amplification. This was supported by the eDNA sequencing of aquarium samples where there was minimal recovery of invertebrate amplicons from tanks known to contain invertebrate species. This suggests the use of MarVer3 in Vertebrate biodiversity surveys would not be limited by potential homology with some invertebrate sequences.

### Performance of MarVer1 and MarVer3 primer sets with environmental samples

We evaluated the performance of MarVer1 and MarVer3 primer sets with water samples collected at the Genoa aquarium, from tanks with known community compositions. Amplicons annotated to species level were recovered for 81.5% and 37% of the 27 resident taxa for MarVer3 and MarVer1 respectively. For MarVer3, the five ‘undetected’ species included *Diplodus cervinus* (Zebra seabream), *Pterygoplichthys gibbiceps* (Armored catfish), two ray species (*Dasiatis americana, Teaniura grabata*), and *Pristis zjisron* (Longcomb sawfish). In the case of *D. cervinus*, amplicons assigned to other non-resident *Diplodus* species were observed, so eDNA from the species may have been present, but annotated as a congeneric. For the four other ‘undetected’ species, there were no other incompletely (above species level) assigned reads at genus, family, or order level which could be attributed to these taxa (for *Pristis zjisron* no matching reference sequence for the MarVer3 region was available in Genbank). These four cases therefore appear to be genuine non-detections. These are all bottom dwelling species, whereas our water samples were collected at the surface, and therefore potentially we did not capture eDNA from these species.

For MarVer1, 9 out of 13 resident species with Genbank reference sequences covering the MarVer1 region were detected successfully. Amplicons correctly assigned to the species level were not observed for the two penguin species, Blackspot seabream (*Pagellus bogaraveo*), and longcomb sawfish, despite reference sequences being available. For the Magellanic penguin, amplicons assigned to the congeneric *Spheniscus demersus* were observed, but no candidate incompletely or misattributed amplicons for the Gentoo penguin (*Pygoscelis papua*) or two fish species were recorded.

For 14 species (51.6% cases), no Genbank reference sequences covering the MarVer1 region were available. Reads assigned to the non-resident grouper *Epinepehelus lanceolatus* were observed, indicating that eDNA from the three resident groupers may have been detected but attributed to a congeneric for which a reference was available. For the remaining 11 species no other candidate amplicons attributable to related taxa were recorded.

Our demultiplexing and annotation pipeline required an amplicon sequence homology of at least 98% with other Molecular Operational Taxonomic Units (MOTUs) and with Genbank reference sequences. Therefore, the lower assignment rate for MarVer1 compared to MarVer3 likely reflects the lower Genbank coverage for the 12S region encompassed by MarVer1. In this case, reducing stringency (e.g. to 95% homology) may increase annotation rates, allowing successful attribution of amplicons to genus and/or family level, but with a requirement to check homology level for individual MOTUs before accepting species level assignments. For both primer sets, annotation success rates would be expected to increase as taxonomic coverage of Genbank reference sequences improves over time. Similarly, while MarVer3 was predicted to potentially recover invertebrate amplicons when allowing for low levels of degeneracy at priming sites, few were observed. Potentially this may also be accounted for by low coverage with reference sequences and the level of stringency applied in the annotation pipeline. The annotation of the few observed invertebrate amplicons from the aquarium samples should also be treated cautiously for the same reasons.

### Optimising locus choice for different eDNA and taxon detection applications

The four loci described in this paper provide investigators with flexible options to target different barcode markers depending on priorities for their study objectives, tailored to requirements for taxonomic breadth, variation and resolution at different taxonomic levels, amplicon size where eDNA degradation is a concern (e.g. Speller et al. 2016), and sensitivity to contamination from human DNA.

Overall the 16S based MarVer3, the largest amplicon product with an expected size of approximately 245bp, has the highest potential taxonomic coverage across all vertebrates, and taxon resolution from species level upwards. Our trial eDNA HTS assay using samples from aquarium tanks demonstrated the locus performed well with environmental samples despite its relatively large amplicon size compared to the other loci. However, the MarVer3 region showed limited variability at nominal species level compared to MarVer1 and 2 amplicons (e.g. in killer whale, *Orcinus orca*, no variability was found among 151 individuals tested, see Table 3). This is consistent with lower rates of evolution in 16S compared to 12S genes, and suggests 12S genes may be more informative for resolving intraspecific differences (see below).

The 12S based MarVer1, Marver2, and Ceto2, offer the advantage of smaller amplicon sizes (approximately 202bp, 89bp, 104bp respectively), which may be a consideration for applications requiring work with more highly degraded eDNA templates (Nichols et al. 2018, Wei et al. 2018).

Here we focused on evaluating taxonomic resolution with marine mammal groups, and found the relative performance of the loci varied across different families. For instance, MarVer1 was more variable than MarVer2 in most Cetacean species, but MarVer2 still had strong taxon resolution for species outside the *Delphinidae* family, and in particular for the *Ziphiidae*. In a stretch of less than 100bp, the MarVer2 region contained more polymorphic sites in the Ziphiidae (5 Genera assessed) than in the numerically and morphologically more diverse Delphinidae Family (14 Genera assessed). This observation is likely due to the older phylogenetic divergence of the Ziphiidae lineages, allowing more time for accumulation of fixed differences between species, compared to the more recently derived Delphinidae family (Nikado et al. 2001). MarVer2 was also more effective at resolving Pinnipeds of the genus *Pusa* compared to MarVer1. However, the incomplete resolution of MarVer2 (and thus Ceto2), within the *Delphinidae* would preclude its use for species level detection studies within the Mediterranean, one of our geographic areas of interest, since it cannot differentiate bottlenose, common and striped dolphins, which are sympatric in this sea. There were also notable performance differences for loci at the interspecific level within Families. For example, MarVer3 was the most polymorphic locus within the 104 spinner dolphin (*Stenella longirostris*) samples, but it was monomorphic within the 151 killer whale (*Orcinus orca*) mitogenome sequences retrieved from GenBank, see Table 3).

Resolution of intraspecific phylogeographic variation is likely to be best attempted with either more variable, or longer target amplicons (e.g. d-loop region; Kunal et al 2013). However, the large Cetacean sequence dataset we evaluated allowed us to test the potential of our loci to identify phylogeographically informative variation, which could be used for simple haplotype clade determination with eDNA for some species (Adams et al. 2019). For instance, within the MarVer1 amplicon in the 151 killer whale mitogenomes (Morin et al. 2010, 2015 and Filatova et al. 2018), some variable sites were private to either the Transient clade or to the AntB and AntC clades identified by the larger dataset.

The preliminary investigation of sequence variation in other large marine vertebrate groups (tuna and sea turtle species), suggested our loci also have potential to be informative for species identification in those taxa. While not assessed directly in this study, the MarVer loci may also prove to be useful as barcode markers for terrestrial vertebrates given taxonomic conservation of the priming sites.

Finally, the high levels of diagnostic variation seen within MarVer loci amplicons offer potential for designing additional species specific nested internal primers (Stoeckle et al. 2018). These might have utility for species focused, non-sequencing based detection applications, such as quantitative PCR, or simple agarose gel based amplicon visualisation when there is limited access to laboratory facilities, or funding. MarVer2 and Ceto2, would be most suited for this in Cetaceans (due to the clustering of variable sites in proximity of one of the priming site), while for Teleost species, MarVer3 would be the best candidate (due to many in-del mutations in fish).

## Conclusions

This paper presents four novel primer sets targeting 12S and 16S vertebrate mitogenome regions, with a particular focus on marine mammals. Using a combination of *‘in silico*’ validation, and application to eDNA samples from aquarium communities with known species composition, we show the loci to have high potential for metabarcoding and eDNA studies targeting marine vertebrates. These primer sets have broader taxonomic coverage and resolution compared to previously developed 12S and 16S primer sets, potentially allowing surveys of complete marine vertebrate communities (including fish, sea turtles, bird and mammals) in single HTS runs, simplifying workflows, reducing costs, and increasing accessibility to a wider range of investigators. They may be applied in any context focusing on resolving vertebrate taxonomic identity, from biodiversity surveys and forensics (e.g. CITES surveillance, or surveys of commercially targeted fish species), through to behavioural ecology studies and supporting conservation of rare or endangered marine vertebrate species.

## FUNDING

No funding was received for this work.

## CONFLICT OF INTEREST

The authors declare that they have no conflict of interest.

## ACKNOWLEDGEMENTS

We thank Fulvio Maffucci for providing marine turtles DNAa samples. Giudo Gnone of the Aquarium of Genoa for allowing and supporting collection of controlled environmental eDNA samples. Antonia Bruno for advises on filtering procedures of environmental samples and Anna Sandionigi for bioinformatic advice.

## AUTHOR CONTRIBUTIONS

E.V. designed primer sets, planned the testing approach compiled the marine mammal guideline to the primers’ use and wrote the manuscript; J.B. performed the *in silico* validation of the novel primer sets; R.L. carried out samples collection, wet lab tests, eDNA amplifications and analysed controlled environment HTS data; S.G. contributed to the design and implementation of the wet lab primer validation, provided support and facilities for HTS analysis at UoL, and contributed to the drafting and editing of the manuscript; I.C. provided HTS services and bioinformatics support; L.C. allowed and supported collection of water samples from tanks of the Aquarium of Genoa structure; A.G. provided useful comments on manuscript; P.G. enthusiastically supported and hosted the research study at UnMB.

---

**Online Resource 1.**
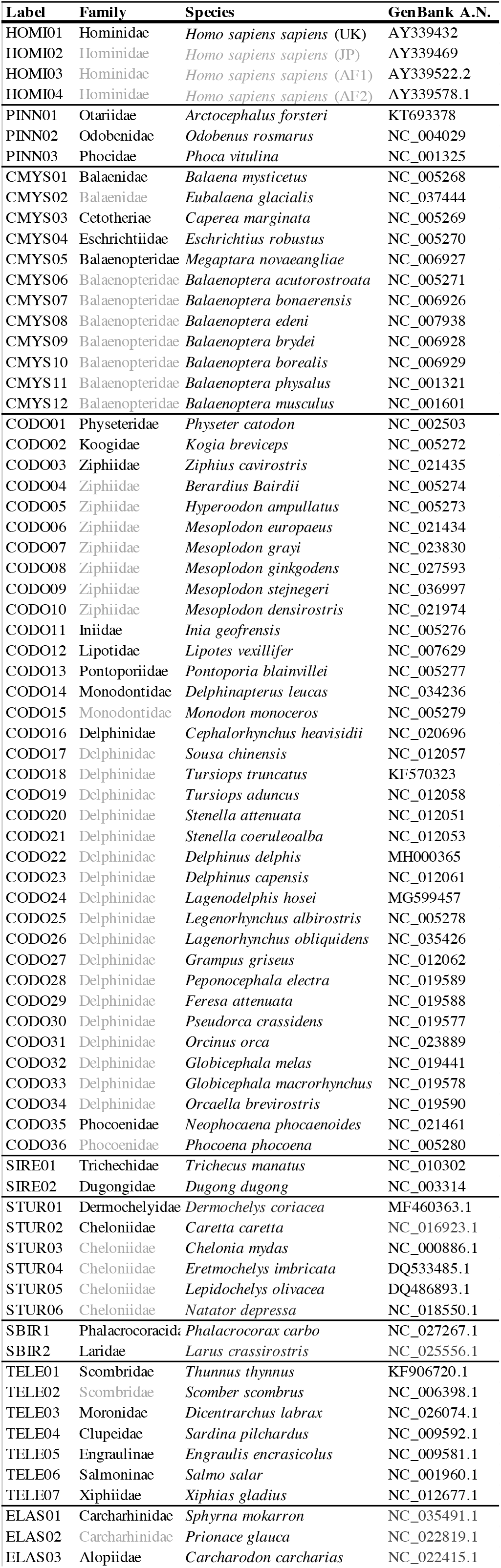
List of 75 taxa used for primer sets’ design.

**Online Resource 2.**
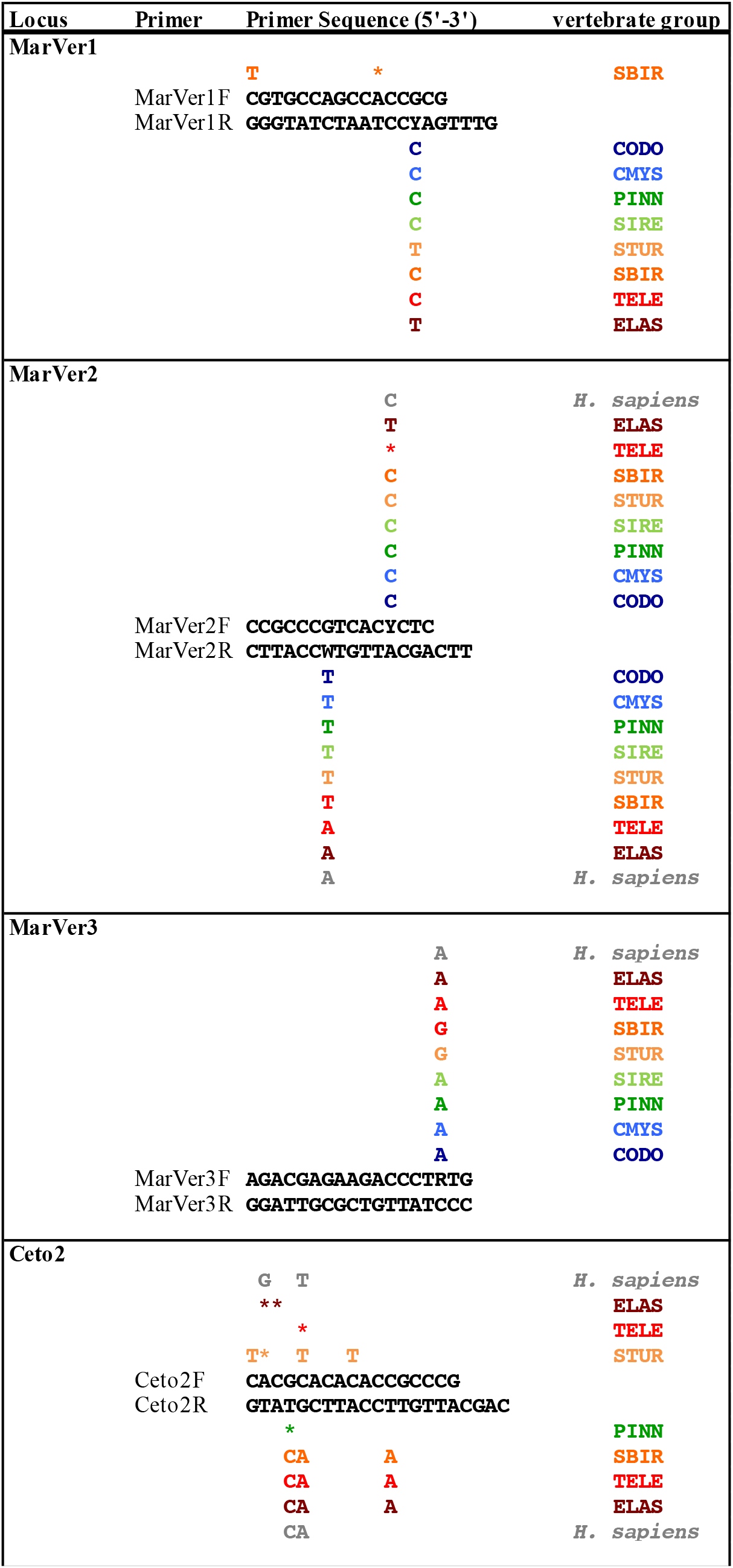
Variable sites at the priming sites of the described loci for each of the eight marine vertebrate groups considered in this study (71 taxa considered): CODO= Cetacean Odontocetes; CMYS= Cetacean Mysticetes; PINN= Pinnipeds; SIRE= Sirenians; STUR= Sea Turtle; SBIR= Sea Birds; TELE= Teleosts or bony fish; ELAS= Elasmobranch or cartilaginous fish. Star signs (*) indicate variable sites within the corresponding vertebrate group.

**Online Resource 3.**
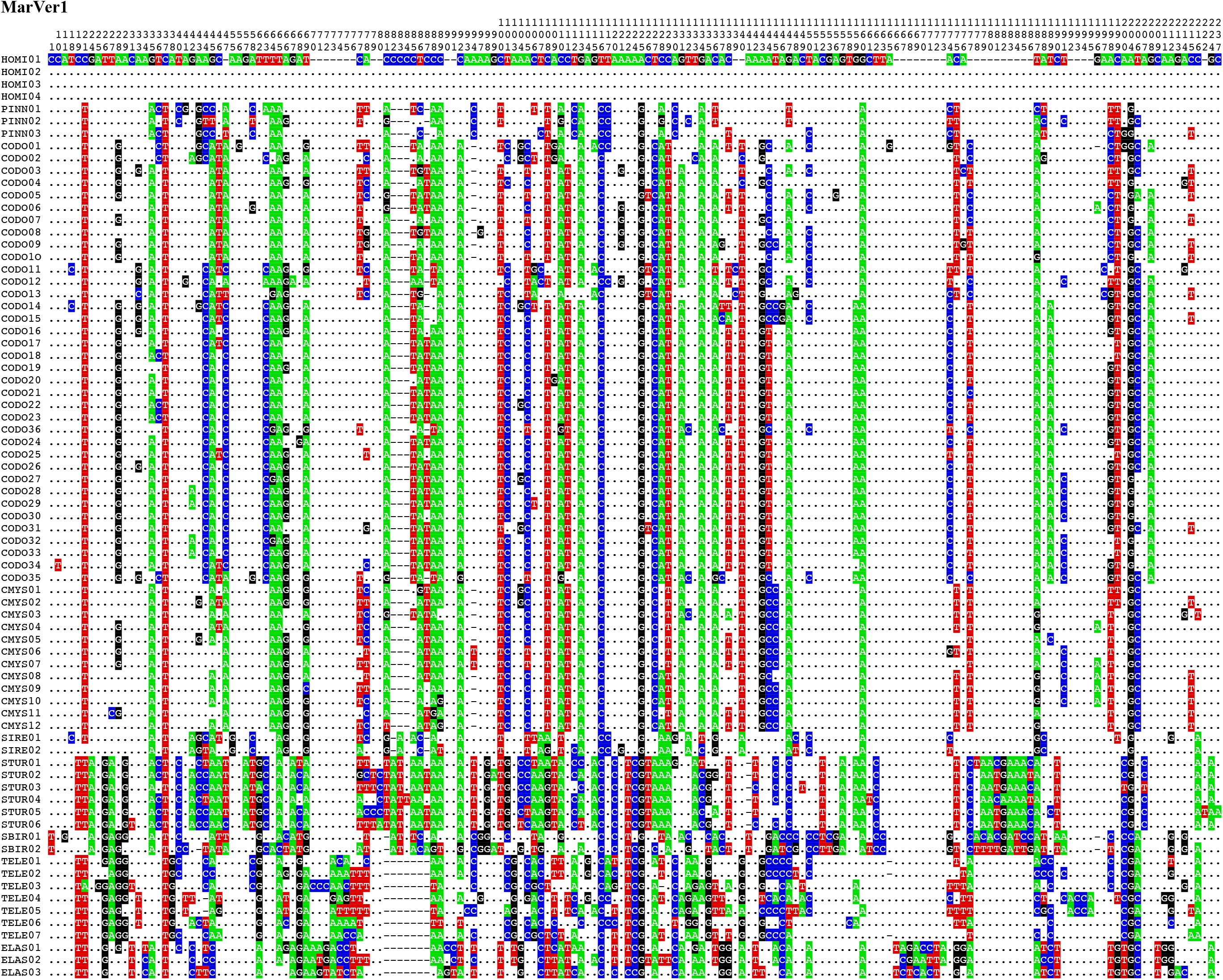
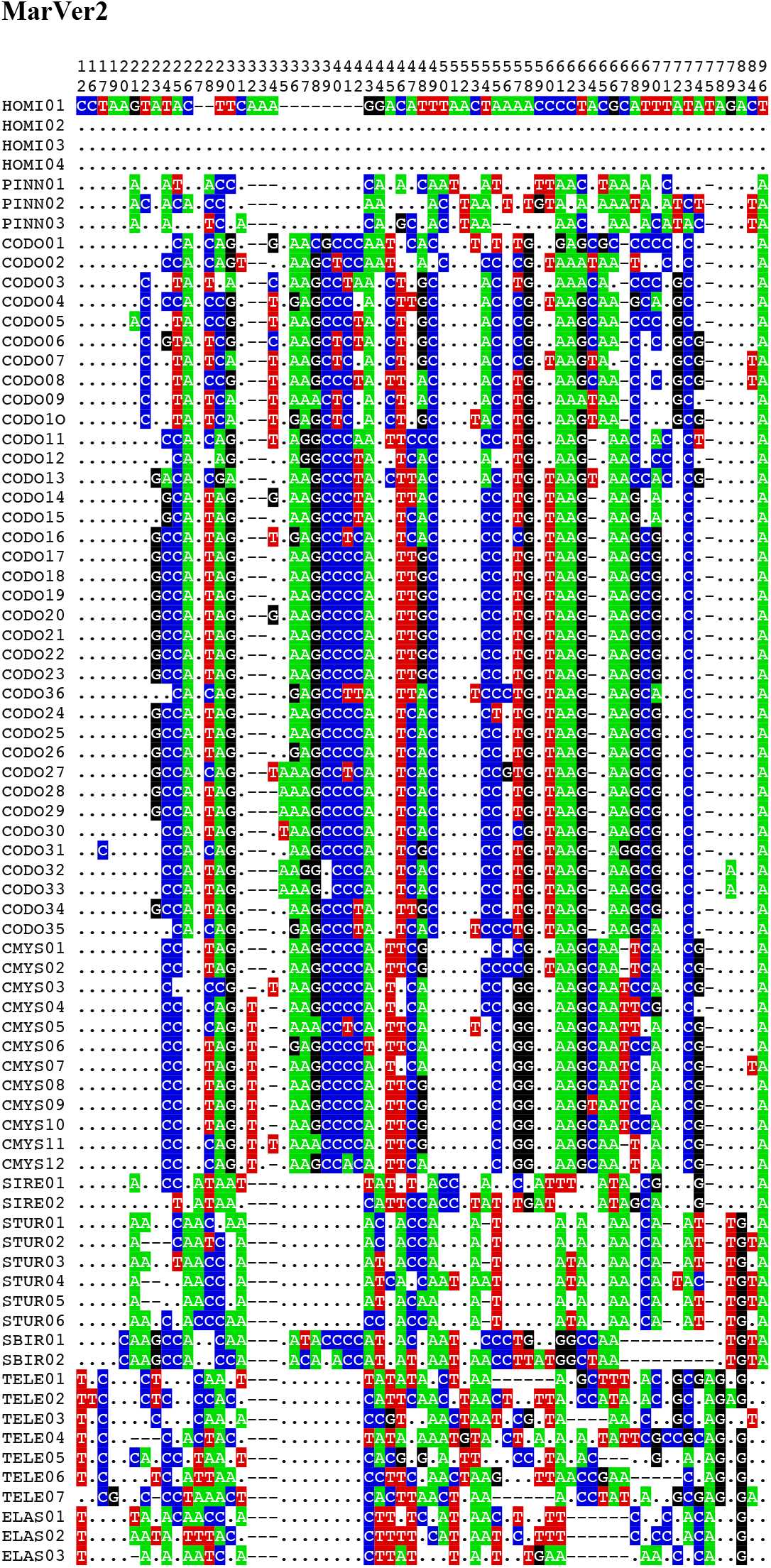
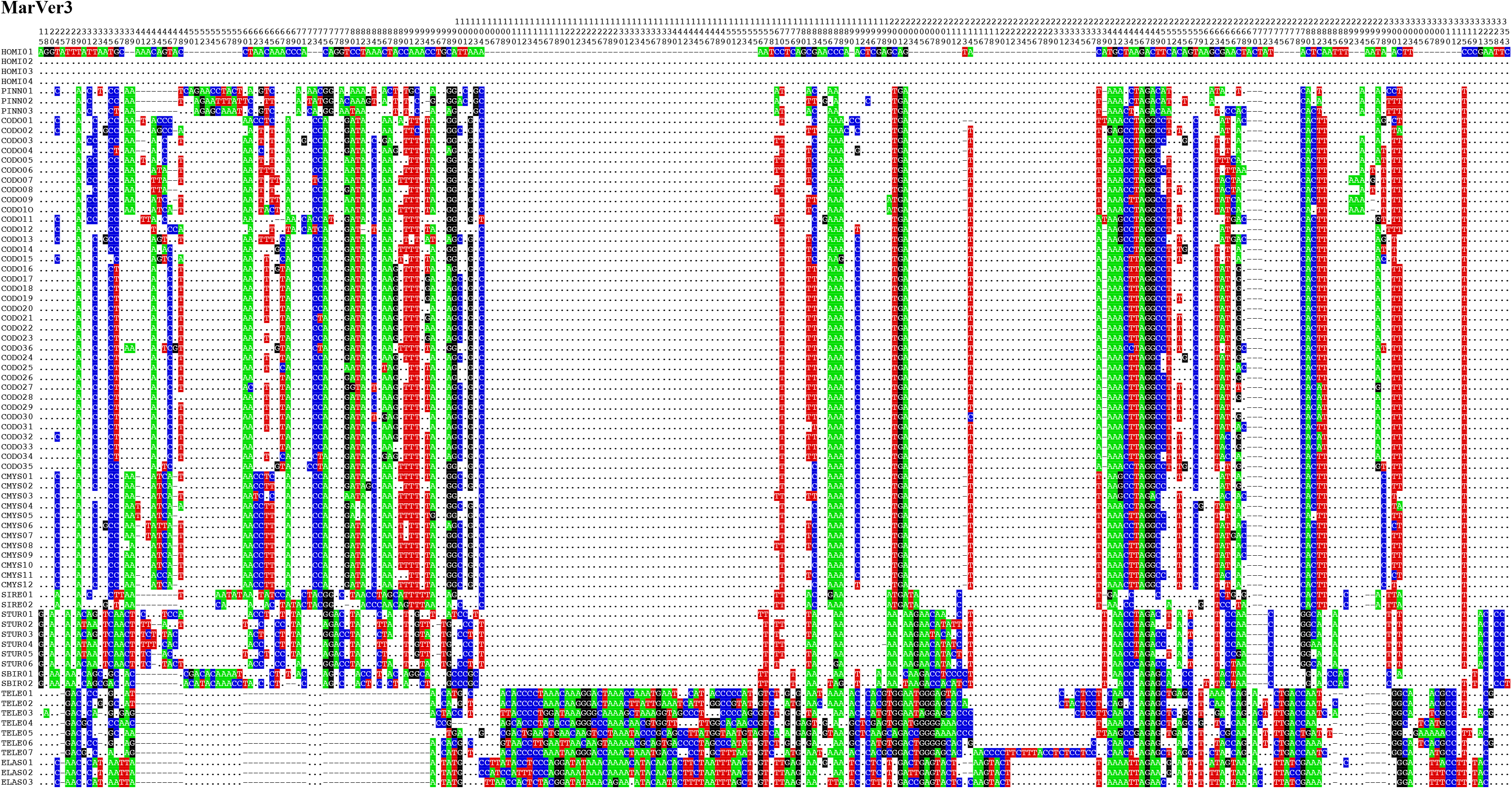
Extraction of variable sites for loci MarVer1, MarVer2, and MarVer3 for the 75 taxa used for primers design.

**Online Resource 4.**
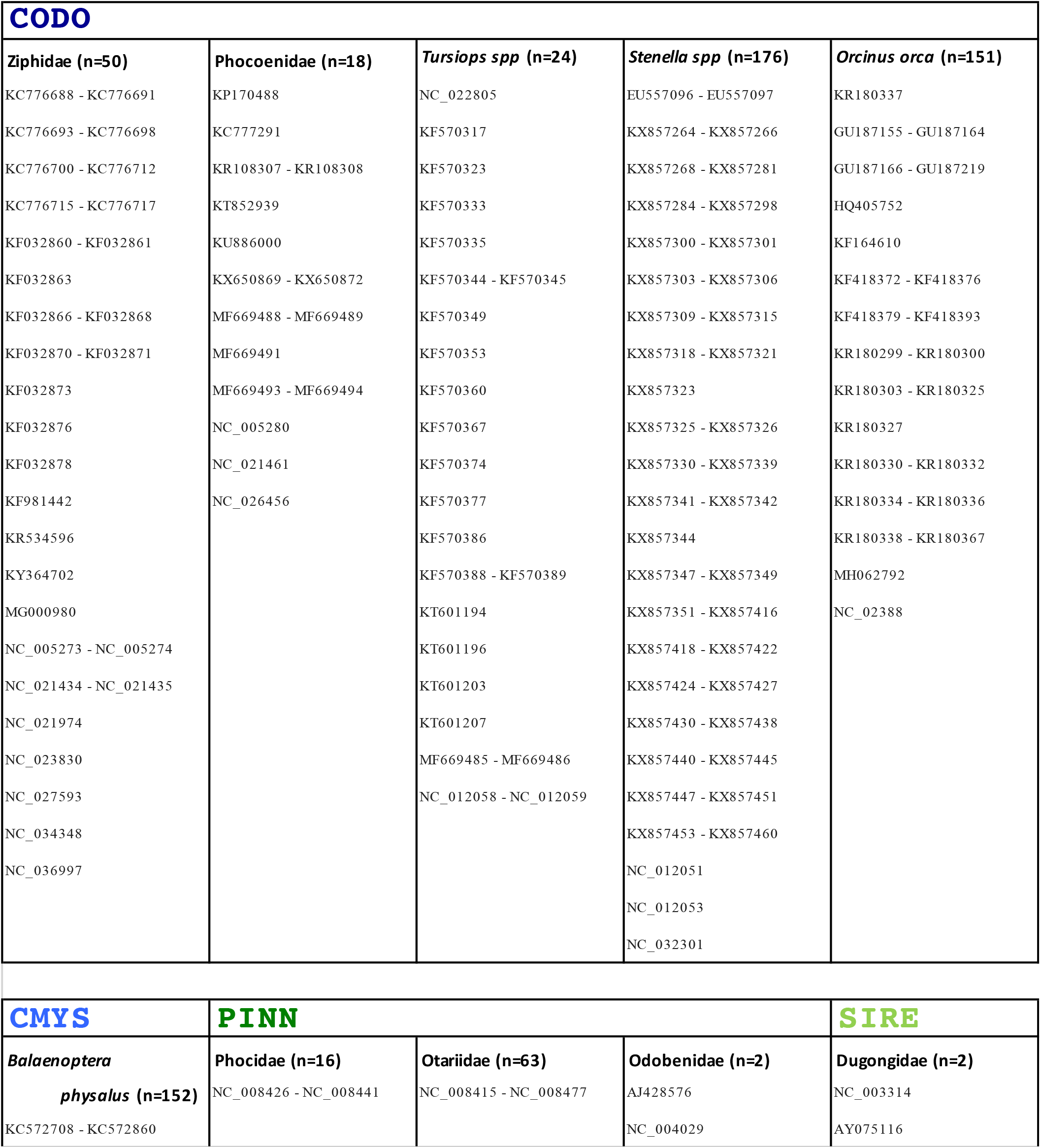
List of 657 GenBank entries used in this study to assess amplicons’ polymorphism at the three presented loci. NB 22 *Pusa capsica* sequences are not shown as unpublished yet.

**Online Resource 5.**
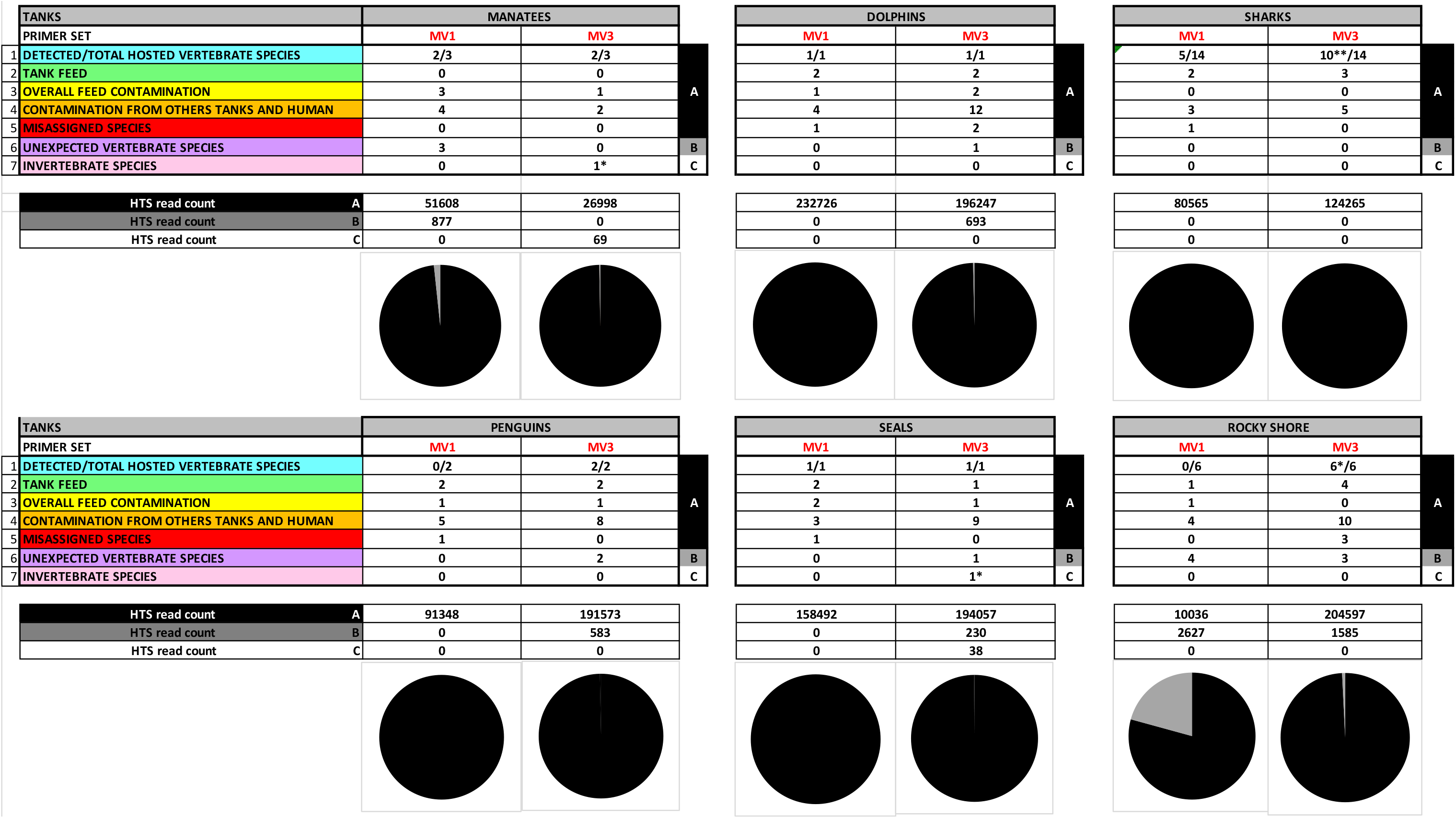
HTS species detection differences between loci MarVer1 and MarVer3, as inferred from controlled environment eDNA samples collected in 6 tanks of the Aquarium of Genoa (July 2018). For each tank, two sets of data are shown: one referring to the number and composition of detected species, where the species scoring more than 100 reads were considered as “detected” (top part of each section), and one showing the number of HTS reads obtained (bottom part of each section, with related pie graphs). Note that all data refer to 6 eDNA-samples replicates collected in each tank. **The top part of each table section shows 7 categories:** 1) “DETECTED/TOTAL vertebrate species” indicates the number of vertebrate species correctly assigned over the total number of species known to be present in the tank as main occupants (“*” indicates two instances -Shark and Rocky shore” tanks-in which one of the species detected was found but with a read count lower than 100); 2) “TANK FEED” indicates the number of molecularly detected species that were not present in the tank as living species, that however were used as food source provided to the tank hosts (thus their DNA traces originating from both hosts’ faeces and food left-overs could be found); 3) “OVERAL FEED CONTAMINATION” indicates the number of molecularly detected species that should not be in that specific sampled tank, but that are found elsewhere in the Aquarium structure, used as food source in other tanks (food buckets and/or operators’ boots or wetsuit are likely to have provided a vehicle for contamination); 4) “CONTAMINATION FROM OTHER TANKS AND HUMANS” indicates the number of molecularly detected species that matched with either species present in other tanks of the Aquarium or humans (NB: the diver in charge of the maintenance of different tanks is the same person, wearing the same wetsuit: often tank hosts -like dolphins- closely interact with him and his wetsuit); 5) “MISASSIGNED SPECIES” groups the molecularly detected species which were not those present in the tank, that however were taxonomically related to them. 6) “UNEXPECTED VERTEBRATE SPECIES” indicate the number of molecularly assigned species whose detection could not be explained. Note that all species falling in these first 6 categories were all vertebrate species (including those falling into the two “food” categories). 7) “INVERTEBRATE SPECIES” indicate the number of molecularly assigned invertebrate species. **The bottom part of each table section shows the number of HTS reads scored in each of the following 3 categories:** A) vertebrate species whose presence (direct or indirect) in form of eDNA traces could be justified (thus grouping together categories 1 to 5 of the above described list) [no artefact] B) vertebrate species whose presence (direct or indirect) in form of eDNA traces could not be explained [possible artefact] C) invertebrate species detection [primer aspecificity] Results are discussed in the main text.

